# Functional Cell Types in the Mouse Superior Colliculus

**DOI:** 10.1101/2022.04.01.486789

**Authors:** Ya-tang Li, Markus Meister

## Abstract

The superior colliculus (SC) represents a major visual processing station in the mammalian brain that receives input from many types of retinal ganglion cells (RGCs). How many parallel channels exist in the SC, and what information does each encode? Here we recorded from mouse superficial SC neurons under a battery of visual stimuli including those used for classification of RGCs. An unsupervised clustering algorithm identified 24 functional types based on their visual responses. They fall into two groups: one that responds similarly to RGCs, and another with more diverse and specialized stimulus selectivity. The second group is dominant at greater depths, consistent with a vertical progression of signal processing in the SC. Cells of the same functional type tend to cluster near each other in anatomical space. Compared to the retina, the visual representation in the SC has lower dimensionality, consistent with a sifting process along the visual pathway.

## 1 Introduction

Parallel processing of information by the brain operates at three different levels: single neurons, cell types, and neural pathways. Processing of visual information at the cell-types level starts at the retina, where the signal is split into ∼15 types of bipolar cells (Shekhar et al., 2016), ∼63 types of amacrine cells (Yan et al., 2020), and ∼40 types of retinal ganglion cells (RGCs) (Roska and Meister, 2014; Sanes and Masland, 2015; Baden et al., 2016). The RGCs send the processed information directly to the superior colliculus (SC), an evolutionarily conserved structure found in all vertebrates (Isa et al., 2021; Basso and May, 2017).

In the rodent, more than 90% of RGCs project to the superficial layer of the SC (Ellis et al., 2016), and each SC neuron receives inputs from about six RGCs (Chandrasekaran et al., 2007). It remains unclear how the information is transformed at this stage and how many cell types exist in the SC. As in other brain areas, a solid classification of cell types in this circuit would support a systematic study of its function (Zeng and Sanes, 2017). By classic criteria of cell morphology and physiology, prior work has distinguished five cell types (Langer and Lund, 1974; May, 2006; Gale and Murphy, 2014) in the retino-recipient superficial layer. Differential expression of molecular markers has been used to describe about ten types (Byun et al., 2016). By contrast, recent work on the primary visual cortex (V1) identified 46 types of neurons based on morphology and electrophysiology (Gouwens et al., 2019). Because the major inputs to the superficial SC are from the retina and V1, we hypothesize that the number of functionally distinct cell types in the SC has been underestimated.

The present work aims to identify functional cell types in the superior colliculus by virtue of their responses to a large set of visual stimuli. These include a panel of stimuli that successfully separated ∼40 types of retinal ganglion cells, confirming many classes previously known from anatomical and molecular criteria (Baden et al., 2016). By two-photon calcium imaging we recorded neuronal responses from the posterior-medial SC of behaving mice while leaving the cortex intact. We classified cell types based on their response to the high-dimensional visual stimulus using unsupervised learning algorithms. We included several transgenic mouse strains that label subsets of SC neurons based on gene expression patterns. The evidence points to ∼24 functional types that come in two major classes: one closely related to retinal responses, the other distinct. We report on the anatomical organization of these functional types, their relation to molecular cell types, and their progression throughout layers of the superficial SC. By comparing the space of visual features encoded in the SC to that in the retina one finds that the superior colliculus already discards substantial information from the retinal output.

## 2 Results

### 2.1 Single-cell imaging reveals diverse neuronal responses to a set of visual stimuli

To investigate the functional diversity of SC neurons, we imaged neuronal calcium responses to a battery of visual stimuli using two-photon microscopy in head-fixed awake mice (Figure 1A). To maintain the integrity of the overlying cortex, one is limited to the posterior-medial SC that corresponds to the upper lateral visual field (Feinberg and Meister, 2015) (Figures 1B and 1C). We recorded more than 5000 neurons from 41 image planes in 16 animals with different genetic backgrounds, including wild-type, Vglut2-Cre, Vgat-Cre, Tac1-Cre, Rorb-Cre, and Ntsr1-Cre mice. In the Cre lines, the calcium indicator was restricted to the neurons expressing the respective transgene.

**Figure 1:**
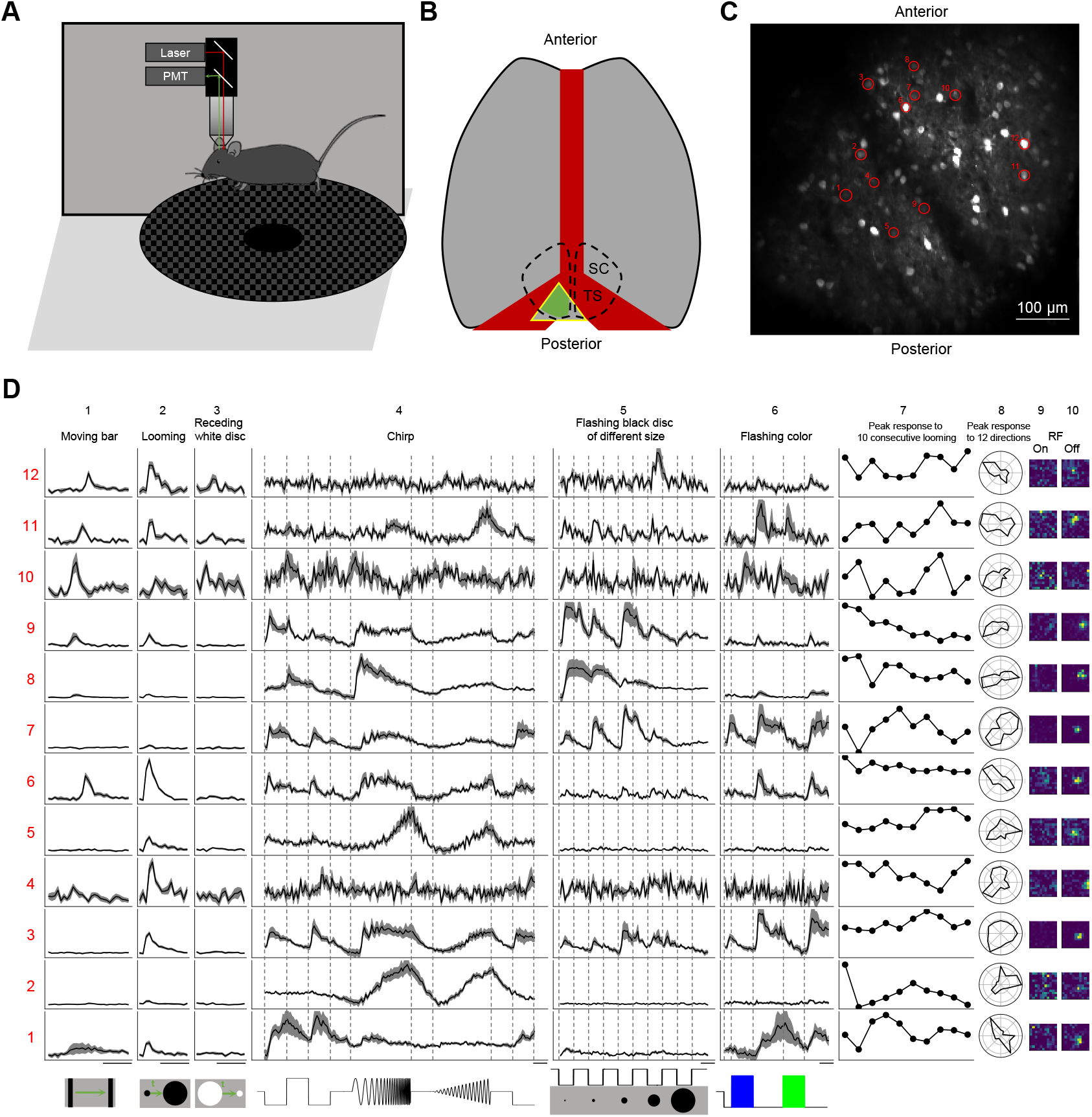
Two-photon imaging reveals diverse visual responses in awake mouse superior colliculus. **A**. Schematic of the experimental setup. Mice were head-fixed and free to run on a circular treadmill. Visual stimuli were presented on a screen. Neuronal calcium activity was imaged using two-photon microscopy. PMT, photomultiplier tube. **B**. Schematic of mouse brain anatomy after insertion of a triangular transparent plug to reveal the posterior-medial part of the superior colliculus underneath the transverse sinus. TS: transverse sinus. **C**. A standard deviation projection of calcium responses to visual stimuli in one field of view. **D**. Response profiles of 12 neurons (rows) marked in **C**. Columns 1-6 are time-varying calcium responses to visual stimuli. Gray shade indicates the standard error across identical trials. Each row is scaled to the maximal response. Scale bars: 2 s. Subsequent columns show processed results: (7) response amplitude to an expanding dark disc on 10 consecutive trials. (8) polar graph of response amplitude to moving bar in 12 directions. (9 and 10) Receptive field profiles mapped with small squares flashing On or Off.

We presented a battery of visual stimuli (Figure 1D) chosen to probe spatio-temporal integration, color-sensitivity, and movement processing (see Section 4.4 for detail). Included was a “chirp” stimulus that modulates the intensity on the cell’s receptive-field over both frequency and amplitude; this was previously employed in the classification of retinal ganglion cells (Baden et al., 2016). Neurons in any given image plane showed robust and diverse responses to these stimuli (Figures 1C-D). The animals were positioned on a circular treadmill but remained stationary during most of the visual stimulation. Because locomotion barely modulates the visual responses of SC neurons (Savier et al., 2019), we did not consider further the effects of movements.

### 2.2 Superficial superior colliculus comprises at least 24 functional cell types

To classify cell types based on their functional properties, we first performed a sparse principal component analysis on the raw response traces (Figure 6A) (Mairal et al., 2009), which led to a 50-dimensional feature vector for each neuron. Then we added 4 designed features that describe different aspects of the response: a habituation index (HI) computed from repeated stimuli, a direction selectivity index (DSI), an orientation selectivity index (OSI), and a motion selectivity index (MSI, see Methods). We focused on 3414 neurons that responded reliably to visual stimuli (signal-to-noise ratio > 0.35, see definition in Methods Eqn 2), and searched for clusters in the 54-dimensional feature space by fitting the data (3414 cells × 54 features) with a Gaussian mixture model, varying the number of clusters in the mixture (Figure 2).

**Figure 2:**
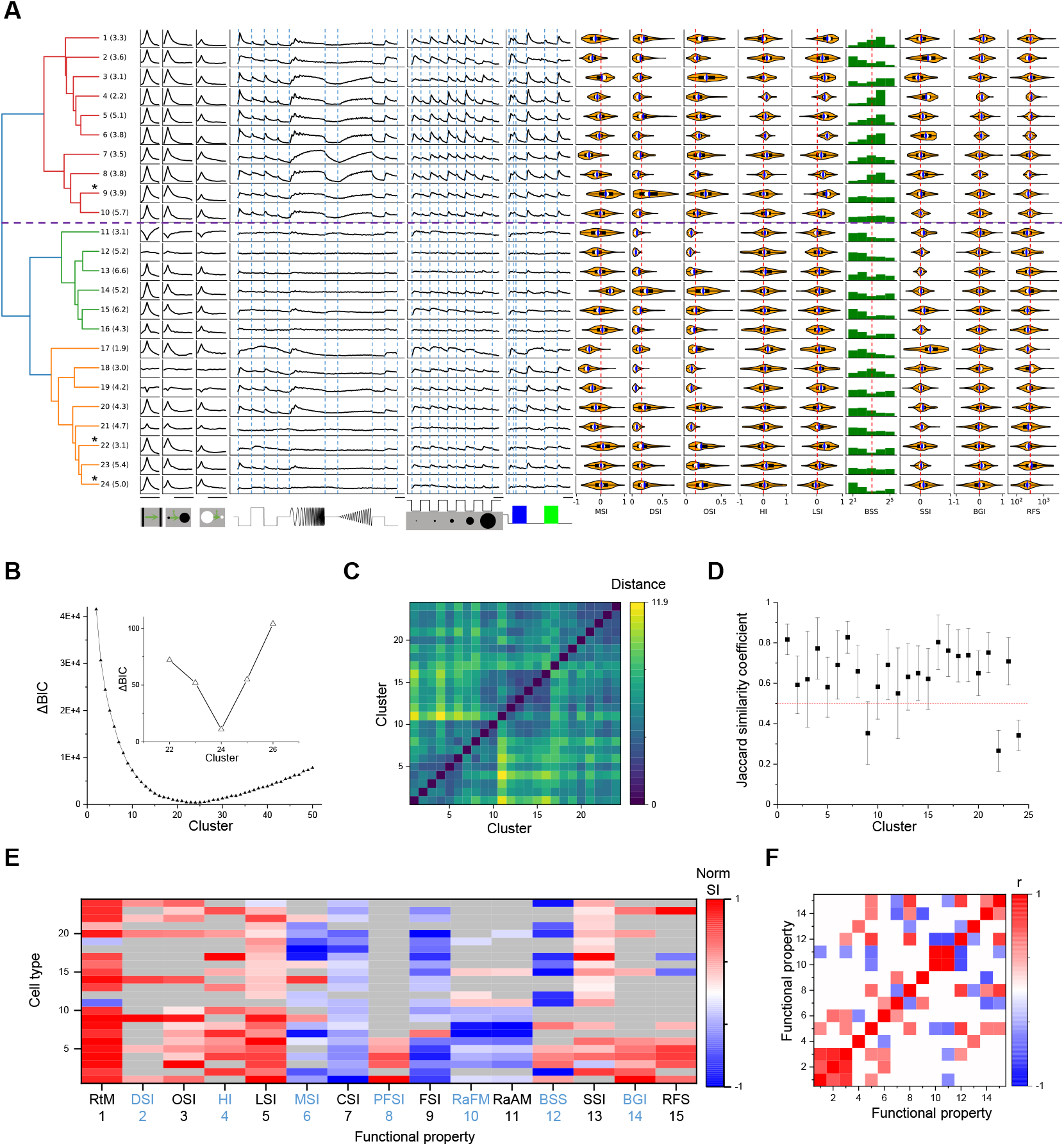
Twenty-four functional cell types in the mouse SC. **A**. Dendrogram of 24 clusters based on their distance in feature space. For each type this shows the average time course of the neural response to the stimulus panel; the vertical scale is identical for all types and stimulus conditions. This is followed by violin plots or histograms of various response indices: motion selectivity index (MSI), direction selectivity index (DSI), orientation selectivity index (OSI), habituation index (HI), looming selectivity index (LSI), best stimulus size (BSS), surround suppression index (SSI), blue green index (BGI), and receptive field size (RFS). Purple dashed line separates groups 1 and 2. Numbers in parentheses indicate percentage of each type in the dataset. Stars mark the unstable clusters with JSC*<* 0.5 (see panel D). Scale bars: 2 s. **B**. Relative Bayesian information criterion (ΔBIC) for Gaussian mixture models with different numbers of clusters. **C**. Euclidean distance between the center of two clusters in the feature space. Clusters are sorted as in the dendrogram (panel A). **D**. Jaccard similarity coefficient (JSC) between the full dataset and subsets (Mean ±SD). **E**. Normalized selectivity index (normalized for each column) of functional properties represented by different cell types (See Methods). SIs that are not significantly different from 0 is shown as grey color. RtM: response to motion; CSI: contrast selectivity index; PFSI: peak-final selectivity index; FSI: frequency selectivity index; RaFM: response after frequency modulation; RaAM: response after amplitude modulation. **F**. Pearson’s correlation coefficients of the representation between pairs of functional properties. Grey color indicates two properties are not significantly correlated.

The quality of each clustering was assessed with the Bayesian information criterion (BIC), which addresses concerns about overfitting by balancing the goodness of fit with generalizability. By this measure the distribution of cells in feature space was best described with a Gaussian mixture of 24 components (Figure 2B), suggesting there are 24 functional types of neurons in the surveyed population. Because the optimum in the BIC curve was rather broad (Figure 2B), and one could make a case for both fewer or more clusters, we followed up by testing the stability of each cluster: We fitted various sub-samples of the data set and assessed how well the resulting clusters correspond to those in the full set, using a number of established statistics (see Methods and Figure 6). It emerged that 3 of the 24 clusters are somewhat unstable (Figure 2C); they are marked as such in Figure 2A. Overall the stability of the cluster definitions matched or exceeded those in related studies of neuronal cell types (Gouwens et al., 2019; Baden et al., 2016). For the purpose of subsequent analysis we will adopt this division into 24 types as suggested by the BIC.

The hierarchical relationship between these functional types is illustrated by the dendrogram in Figure 2A, which is based on the distances in feature space between cluster centers. The first branching of the dendrogram splits the types into two groups (Figures 2A, 7, and 8), which we will call Group 1 (types 1-10) and Group 2 (types 11-24). Group 1 further splits into Group 1a (types 1-6) and 1b (types 7-10).

What are the distinguishing features in their visual responses? Figure 2E distills the responses to the stimulus palette into 15 indices (see Methods) that help to characterize each type. As a rule, almost all the types are sensitive to moving stimuli, like the traveling dark bar and the expanding dark disc (RtM in Figure 2E). For many types, these were the stimuli that elicited the strongest response.

Group 1 (types 1-10) is distinct from Group 2 (11-24) in that it responds more strongly to the chirp stimuli (flashes and sinusoid modulations in Figure 2A). All types in Group 1 prefer the expanding dark disc over the receding white disc. Almost all these types are excited by both On and Off stimuli in the receptive field. Also they prefer large spots to small spots, type 2 being a notable exception. Within Group 1, types 1-6 (Group 1a) are distinct from types 7-10 (Group 1b) in their response to sinusoid flicker: Group 1a prefers the low frequencies, whereas Group 1b responds over a wider range. Type 7 in particular rejects the low flicker frequencies.

In Group 2, the response to chirp stimuli is generally weak compared to the moving stimuli. These types respond well to small spots, unlike the Group 1 types. Other response properties in Group 2 are more diverse, and some of these types have been noted previously. For example, Types 11 and 19 stand out in that moving stimuli suppress their activity (Ito et al., 2017). Several types (11, 15, 17) are suppressed by the sinusoid flicker (Ito et al., 2017); 11 and 15 also show rebound after cessation of that stimulus. Type 14 responds strongly to moving stimuli but hardly at all to the entire chirp. Type 18 is remarkably insensitive to any moving stimulus.

Some of these features of the visual response were highly correlated with each other, in that they covaried in the same or opposite directions across types (Figure 2F). For example, direction selectivity (DSI) and orientation selectivity (OSI) tend to be strong in the same cell type. Another strong correlation exists between the preferred stimulus size (BSS) and the response during recovery from the frequency and amplitude chirps (RaFM and RaAM). Much of that correlation traces to a difference in these properties between types in Groups 1 and 2 (Figure 2E).

### 2.3 Neurons of the same type cluster in space

In the retina, ganglion cells come in ∼40 different types (Sanes and Masland, 2015), and they tile the surface in a so-called mosaic arrangement. Neurons of the same type are spaced at regular distances from each other, as though they experienced some sort of repulsion. Neurons of different types are distributed more or less independently (Roy et al., 2021); therefore a ganglion cell’s nearest neighbor is almost always of a different type. The presumed purpose of this arrangement is to ensure uniform coverage such that every location in the visual field has access to each of the types of retinal ganglion cell. Because the retina projects directly to the SC, we investigated whether neurons there are also organized for uniform coverage.

Figure 3A illustrates SC neurons in a single image plane, labeled according to functional type (for more examples see also Figure 9A). Several features are immediately apparent. First, cells of a given type do not repel each other; in fact, the nearest neighbor is often a neuron of the same type. Second, the types don’t cover space uniformly. Some types are segregated from each other (e.g., 7 and 21) whereas others overlap in space (e.g. 14 and 24).

**Figure 3:**
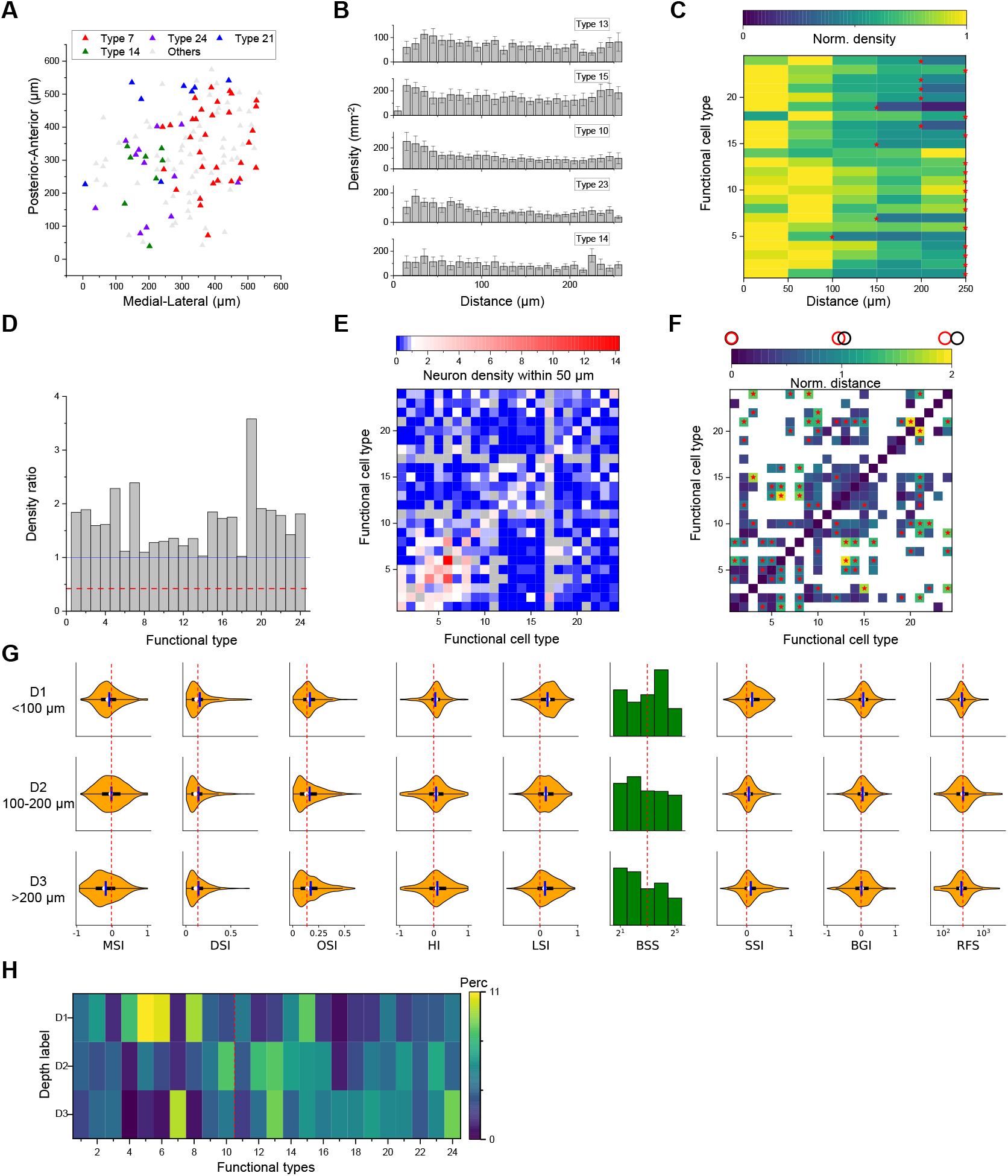
Spatial organization of functional cell types. **A**. Anatomical locations of four types of neurons from one sample recording. **B**. Averaged DRP of five example types for all imaging planes. Error bars denote SEM. **C**. Density of neurons of a given functional type at various distances from a neuron of the same type. Red stars mark the smallest radius within which the neuron density is larger than half of the peak density. **D**. The ratio between density of 0-0.5 RF size and density of 0.5-1 RF size for each type in the averaged DRP. The red dash line is the value for W3 RGC estimated from (Zhang et al., 2012). **E**. Density of neurons in different functional types (columns) within 50 *µ*m of a given neuron whose type is indicated by the row. Note the largest density is the same type except types 8, 11 and 16. Grey color indicates insufficient data for those pairs. **F**. Normalized anatomical distance between neurons in any two functional types. Red stars indicate significant non-overlapping (p<0.05, bootstrap analysis). White color indicates no insufficient data for those pairs. **G**. Functional properties of neurons in three ranges of depth. Display as in Figure 2A. **H**. The percentage of functional types in each depth range. The vertical red dashed line separates types 1-10 and types 11-24.

To pursue these spatial arrangements in greater detail we computed for each functional type the spatial auto-correlation function (also called “density recovery profile”): This is the average density of neurons plotted as a function of distance from another neuron of the same type (Zhang et al., 2012; Rodieck, 1991). When applied to retinal ganglion cells, this function shows a pronounced hole of zero density at short distances. Here, instead, the density remains high down to a distance of 10 µm, which is the typical diameter of a soma (Figure 3B). In fact for several of these SC types the density is highest just 1-2 cell diameters away.

Over larger distances, the autocorrelation function decays gradually (Figure 3C). It drops to half the peak density at a radius of 150-250 µm. This suggests that neurons of a given type form patches of 300-500 µm diameter. Note this accords with a similar patchy organization found previously for certain functional parameters, like the preferred orientation (Feinberg and Meister, 2015) or preferred direction of motion (de Malmazet et al., 2018; Li et al., 2020).

We considered a potential source of error that could give the mistaken appearance of a patchy organization: A functional type might fortuitously be limited to just one recording session, owing to some peculiarity of that animal subject, and so would appear only in the visual field covered during that session. This is not the case: Figure 10 shows that different recording sessions confirm the same functional type. Furthermore, a single recording session reveals separate patches of different types (Figure 3A).

An important scale by which to judge this spatial organization is the size of the receptive field (RF). In the retina, the mosaic arrangement spaces the ganglion cells about one RF diameter apart, so that the RFs of neurons from the same type have little overlap. In the SC that is clearly not the case. From the autocorrelation functions (Figure 3B) we compared the average density of cells within 0.5 RF diameters to the density at 0.5-1.0 RF diameters. In the SC, that ratio is greater than 1 for all functional types and often close to 2 (Figure 3D). By comparison, for the W3 type of retinal ganglion cell (Zhang et al., 2012) that ratio is only 0.4.

In the retina, the arrangement of one functional type is almost entirely independent of the other types (Wassle et al., 1981) (but see Roy et al., 2021). In the SC we found a strong anticorrelation: Within 50 µm of a given cell, the cells of the same type occurred at greater-than-average density, but cells of the other types at lower density (Figure 3E). One might say that each functional type tends to displace the others. To illustrate this further, we calculated the normalized distance between all pairs of cell types (Figure 3F). Twenty-one types show significant spatial separation from at least one other cluster (Figures 9C).

The functional properties of neurons also varied along the depth axis of the superior colliculus. The individual response parameters changed in only subtle ways; for example one can find more neurons with larger receptive fields at greater depth but on average deeper neurons preferred smaller spot stimuli. However, at the level of identified types (which takes many response parameters into account) the differences were more pronounced. In particular, neurons in the upper 100 *µ*m were composed primarily of types 1-10 (Group 1) while neurons deeper than 100 *µ*m belonged primarily to types 11-24 (Group 2, Figures 3H, 9E, p=0.0039, one-way chi-square test).

### 2.4 Genetically labeled populations comprise multiple functional types

Several experiments focused on superior colliculus neurons with specific molecular identity, taking advantage of existing mouse Cre-lines (Figure 4). The Vgat-Cre and Vglut-Cre lines respectively label GABAergic and glutamatergic neurons; Rorb-Cre and Tac1-Cre label populations of neurons stratified in the superficial SC (Byun et al., 2016; Harris et al., 2014); finally Ntsr1-Cre labels a morphological type: the wide-field cells (Gale and Murphy, 2014).

**Figure 4:**
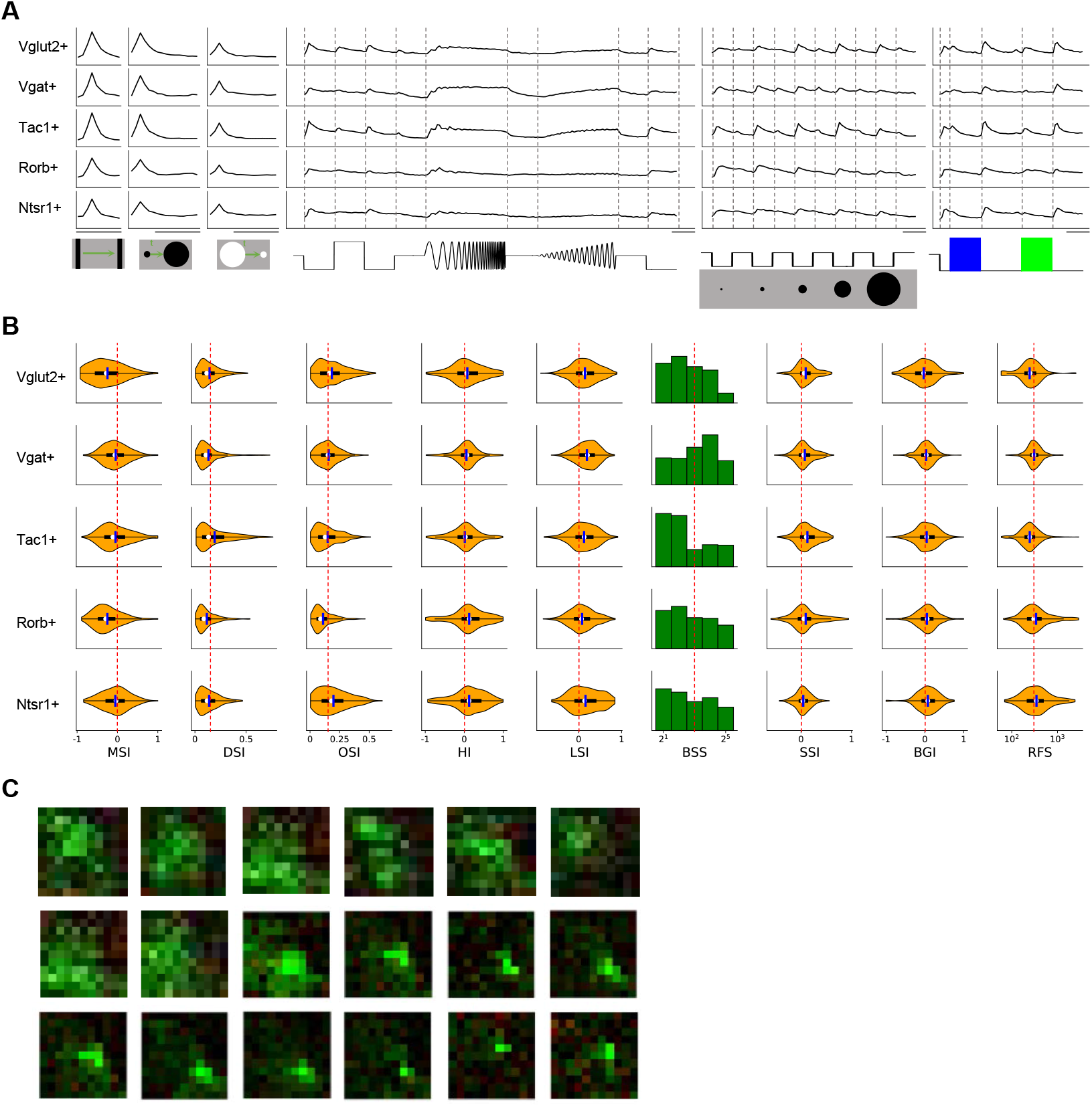
Functional properties in genetically labeled populations. **A**. Average response to the chirp stimulus of five genetically labelled cell types. Display as in Figure 2A; scale bars: 2 s. **B**. Functional properties of the genetically labelled types. **C**. Example receptive fields of Ntsr1+ neurons. Green color indicates the Off subfield and red color indicates the On subfield.

Some systematic differences between these genetically labeled populations are apparent already from the chirp responses (Figure 4A). For example the Rorb+ and Tac1+ neurons are dominated by Off-type responses, whereas ON-Off responses are the rule in the other populations. Excitatory and inhibitory neurons also differ at the population level: Vglut+ neurons prefer flashed over moving stimuli and spots of small size, whereas Vgat+ neurons respond equally to flashed and moving stimuli and prefer large-size spots (Figure 4B). Tac1+ neurons have the highest direction selectivity and small receptive fields, whereas Ntsr1+ neurons show strong orientation selectivity. We refrain from analyzing the fine-grained distribution of the 24 functional types in each genetic population, because the experiments did not cover every combination of genetic label and depth and visual field location, leading to possible confounds.

Prior reports on the Ntsr1+ neurons suggest that they are of the distinctive wide-field anatomical type, with a broad dendritic fan (Major et al., 2000), and that they have the largest receptive fields in the superficial SC (Gale and Murphy, 2014). In fact, we did find very large receptive fields among Ntsr1+ neurons, but surprisingly also many small ones (Figure 4C). Notably another recent study also reported a large variation of receptive field sizes in this line (Hoy et al., 2019). Perhaps some of the Ntsr1+ neurons have the wide-field morphology but are dominated by just one or a few dendritic inputs.

### 2.5 Transformations from retina to superior colliculus

The superficial layers of the superior colliculus receive direct input from most types of retinal ganglion cells. How is visual information transformed as the superior colliculus processes these signals further? To address this directly, our experimental design included a precise copy of the stimuli used previously to classify and distinguish functional types of retinal ganglion cells (Baden et al., 2016). Here we compare the known retinal ganglion cell response types to those we identified in superior colliculus.

As a first-order analysis, we fitted the chirp and color responses of each SC type with a linear combination of the responses of RGC types (Figures 5A, 11A-D). Under the null hypothesis where each SC type is dominated by one RGC type, this should yield fit weights only along the diagonal. Instead, the optimal fit mixes contributions from many RGC types. While these coefficients cannot be interpreted as representing synaptic connectivity, they nonetheless indicate a broad mixing of retinal inputs at the level of superior colliculus.

**Figure 5:**
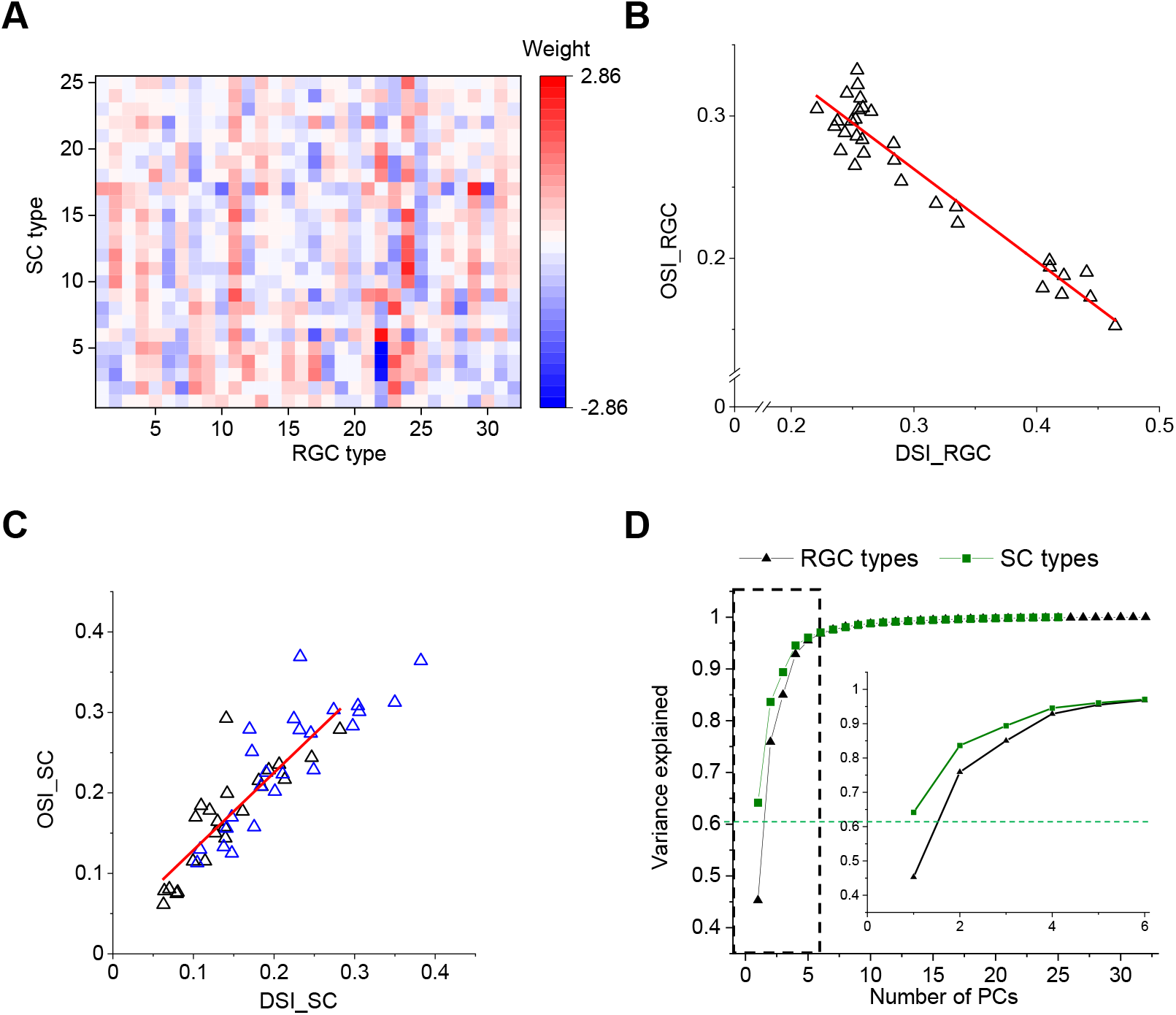
Comparison with functional RGC types. **A**. Weights of different RGC types on each SC type (see Methods). Green dashed line separates types 1-10 and types 11-24. **B**. Plot of OSI versus DSI for 32 functional RGCs (Data are from (Baden et al., 2016)). **C**. Plot of OSI versus DSI for 24 functional types in the SC. Blue triangles denotes OSI and DSI calculated using the same algorithm in (Baden et al., 2016). **D**. PCA on the visual responses of SC and RGC types. Inset enlarges the part in the dashed rectangle.

A number of interesting features emerge. First, some RGC types have universally excitatory (types 4 and 11) or universally inhibitory (types 1, 6 and 20) effects on almost all the SC responses. These RGC types represent a diversity of functional properties. Second, for many RGC types the sign of their contribution changes between the SC types of Group 1 (types 1-10) and Group 2 (11-24). Recall that these groups represent the major division in the dendrogram of SC types (Figure 2A).

Another substantial transformation from retina to colliculus occurs in the representation of orientation and direction for moving stimuli. In the retina, different ganglion cell types are used to encode the direction vs the orientation of bar stimuli (Figures 5B, 11E). By comparison, in the colliculus it appears that the same cells are selective for orientation and for direction (Figures 5C, 11F-I). Similarly, the retina contains many ganglion cell types with purely On-type or Off-type responses (Wang et al., 2010; De Franceschi and Solomon, 2018). By comparison, at the level of the colliculus those two pathways are largely combined, and most functional types are of the On-Off type with a bias towards Off responses (Figure 2A). These examples suggest a reduction in diversity during transformation from retina to colliculus. To test this more globally, we subjected the entire set of chirp-color responses to a principal component analysis, and compared the results from RGC types to those from SC types (Figure 5D). In the superior colliculus, the variance in the visual responses can be explained by fewer principal components than in the retina. This suggests a reduction in diversity from retina to colliculus, as might be expected from selective visual filtering of features essential to guide the animal’s behavior.

## 3 Discussion

Parallel pathways are central to the architecture of biological vision. The visual pathway forks already at the photoreceptor synapse – into ON and OFF representations – and by the time the signal emerges from the eye it has been split into about 40 channels. Each of these pathways is carried by a type of retinal ganglion cell, tuned to certain spatio-temporal features of the visual input, and its neural population covers the entire visual field. How these parallel signals are combined and elaborated in subsequent neural stages of the visual system to sustain the behavioral needs of the animal remains a fundamental question for vision science. Here we sought to follow these parallel pathways into the superior colliculus, the most important retinal projection target in rodents. The approach was to survey visual response properties across many neurons in the superficial layers of the SC, using precisely those stimuli that had been instrumental in defining the parallel pathways at the retinal output.

### 3.1 Main findings

Based on their responses to a broad set of visual stimuli, neurons in the upper SC fall into 24 functional types (Figure 2). At a coarse level, the types form two groups, depending on whether they respond well to simple chirp stimuli (Group 1) or not (Group 2) (Figure 2A). Cells in Group 1 dominate the most superficial regions (<100 um depth), those in Group 2 the deeper levels (Figure 3H). Unlike in the retina – where cells of the same type are spaced laterally at a regular distance – in the SC cells of the same type seem to attract each other. In fact many types appear in clusters, about 300-500 µm in size (Figure 3). Putative excitatory and inhibitory neurons differ in functional properties: On average the inhibitory neurons have a greater preference for large stimuli and for stimulus motion (Figure 4). Compared to retinal ganglion cells, none of the SC types match an RGC type exactly; instead different retinal sensitivities get combined at the level of SC (Figure 5). Overall, the visual representation in the superficial SC is somewhat less diverse than in the retina (Figure 5D).

### 3.2 Relation to prior work

Although the superior colliculus receives more than 40 parallel channels of input from the retina and the visual cortex, prior work classified only 4-5 types of neurons in the SC (Wang et al., 2010; Gale and Murphy, 2014; De Franceschi and Solomon, 2018). The greater diversity of response types in the present study results in part from a different conceptual approach: Instead of defining cell types based on morphological and electrophysiological criteria (Wang et al., 2010; Gale and Murphy, 2014), we focused on visual response properties. This method acknowledges that neurons with the same shape do not necessarily perform the same function in the circuit. Indeed prior work had found that even among neurons with the distinctive wide-field shape there is a great range of response properties (Hoy et al., 2019).

Another supporting factor is the much greater number of neurons in the present survey, which allows finer distinctions and statistical assessment of robustness of the functional cell typing. Further, much of the prior work was conducted under anesthesia, which adversely affects visual selectivity in the superior colliculus (De Franceschi and Solomon, 2018), and thus lowers the number of distinguishable cell types.

At the same time, some of the 24 functional types described here are also evident in prior studies: Types 11 and 15 that show rebounded responses after the frequency-modulated flash are similar to the suppressed-by-contrast cell type (Ito et al., 2017). Types 9 and 14 showing strong orientation and direction selectivity, as reported in prior work (Inayat et al., 2015; De Franceschi and Solomon, 2018).

### 3.3 Implications for visual processing

Regarding the fate of parallel pathways emerging from the retina, one can contemplate two opposing hypotheses: (1) Further divergence of pathways. In this picture downstream circuits split the visual representation further, yielding more functional types of neurons. Each type responds sparsely only when its favorite feature occurs. (2) Convergence of parallel pathways. In this version downstream circuits begin to narrow the visual representation, computing a few variables from the scene that are useful for controlling behavior, while discarding those bits of information that aren’t needed. The present observations favor the second hypothesis, for the following reasons:

First, there are fewer recognizable types in this survey of SC responses (24) than there are in the retinal ganglion cell layer (40) under the same tests. Some comments are in order regarding this comparison. In the retina, the number of RGC types is supported by a convergence of functional, molecular, and anatomical criteria (Sanes and Masland, 2015), leaving little ambiguity. By comparison, the proposed number of 24 functional types in SC is open to debate. The statistical criterion we used (Figure2b) is not the only option, and one can envision a coarser split into fewer types. Furthermore, the cell types reported here, as well as those in previous studies (Wang et al., 2010; Gale and Murphy, 2014; De Franceschi and Solomon, 2018), may well lie at different levels of the synaptic network, whereas the RGCs all lie in the same network layer of parallel representation. All these biases point in the same direction: There are very likely fewer SC types than RGC types. Second, certain features of the visual stimulus that were separated into different retinal cell types appear combined within the superior colliculus. Notably this is the case for the representation of orientation and motion direction (Figure 5C) and for that of On- and Off-signals (Figure 2A). Third, a global measure of dimensionality in the neural representation is smaller among SC types than RGC types (Figure 5D).

All this suggests that the functional diversity in the visual pathway may be greatest at the retinal output. Subsequent circuits may act primarily as a switchboard to distribute these RGC signals to different brain targets. For example, a recent study traced the projections from SC to two different target areas, and found that the corresponding SC cells combine inputs from different subsets of retinal ganglion cells (Reinhard et al., 2019). It will be interesting to explore how these projection-defined types correspond to the functional types reported here and their putative RGC complements (Figure 5A).

Another output target from the superficial SC are the deeper layers, where visual information is combined with other sensory pathways as well as motor signals. These are out of reach for effective optical recording but can be accessed by electrodes. There one encounters response properties not seen in the retina, foremost among them a pronounced habituation to repeated stimuli (Dräger and Hubel, 1975; Horn and Hill, 1966; Lee et al., 2020). Also, the range of response properties narrows systematically with depth, reflecting an increased selectivity for behaviorally-relevant visual features (Lee et al., 2020). Similarly, it is possible that other regions of the SC, outside the posteriormedial sector studied here, contain a different restricted complement of response types.

### 3.4 The spatial organization of functional cell types

The patchy organization of functional cell types (Figure 3) extends the results reported previously regarding specific neuronal response properties, notably the preferred orientation (Feinberg and Meister, 2015; Ahmadlou and Heimel, 2015) or movement direction (Li et al., 2020; de Malmazet et al., 2018). A common theme to all these reports is that the rules of visual processing seem to vary across broad regions of the visual field (for a dissenting report see (Chen et al., 2021)). In the present study we found that different regions of the visual field, measuring ∼ 20-40 degrees across, are covered by a different complement of functional types. Our recordings were limited to the posterior-medial portion of the SC (upper temporal visual field), and thus the global organization of these functional types remains unclear. Also, it is possible that additional functional types will emerge in other parts of the visual field.

It is important to contrast this organization with inhomogeneities found elsewhere in the visual system. Among retinal ganglion cells (RGCs), the mosaic arrangement of neurons within a type guarantees that they are properly spaced, such that each point in the visual field is handled by at least one such ganglion cell. Within a given RGC type, the response properties may vary gradually across the visual field, typically in a naso-temporal or dorso-ventral gradient (Bleckert et al., 2014; Warwick et al., 2018). This is very different from the regional specializations encountered in the SC. In the primary visual cortex of large animals one often finds that functional types are organized in patches or stripes. However, the scale of this functional anatomy is finer than the receptive field size: This guarantees again that each point in the visual field is handled by neurons of every type (Blasdel and Campbell, 2001). By contrast, in the SC the observed patches are 10-20 times larger than receptive fields, which implies a regional specialization of certain visual processes. The reasons for this specialization remain unclear. One can certainly invoke ecological arguments for treating the upper and lower visual fields differently, but on the scale of 30 degrees the purpose is less obvious.

Future work may also inspect the downstream effects of this patchy organization. One hypothesis is that a given functional type gathers visual information distributed to one of the many downstream visual centers. If so, then a retrograde tracing from one of the SC’s target regions should cover only a patch of the visual field. The rapid progress in connectomic and transcriptomic methods for mapping cell types promises further insights into this unusual functional organization.

## 4 Materials and Methods

### 4.1 Animal

Laboratory mice of both sexes were used at age 2-4 months. The strains were C57BL/6J (wild-type), Vglut2-ires-Cre (The Jackson Laboratory, stock No: 028863), Vgat-ires-Cre (The Jackson Laboratory, stock No: 028862), Tac1-IRES2-Cre-D (The Jackson Laboratory, stock No: 021877) (Harris et al., 2014), Rorb-IRES2-Cre-D (The Jackson Laboratory, stock No: 023526) (Harris et al., 2014), and Ntsr1–GN209–Cre (Gerfen et al., 2013). All animal procedures were performed according to relevant guidelines and approved by the Caltech IACUC.

### 4.2 Viral injection

We injected adeno-associated virus (AAV) expressing non-floxed GCaMP6 (AAV2/1.hSyn1.GCaMP6f.WPRE.SV40) into the SC of wild-type mice (C57BL/6J), and AAV expressing floxed GCaMP6 (AAV1.Syn.Flex.GCaMP6f.WPRE.SV40) into the SC of Vglut2-ires-Cre, Vgat-ires-Cre, Tac1-IRES2-Cre-D, Rorb-IRES2-Cre-D, and Ntsr1–GN209–Cre mice. After 2–3 weeks, we implanted a cranial window coupled to a transparent silicone plug that rested on the surface of the SC and exposed its posterior-medial portion. This portion of the SC corresponds to the upper-temporal part of the visual field. The optics remained clear for several months, which enabled long-term monitoring of the same neurons. Two-photon microscopy was used to image calcium signals in the SC of head-fixed awake mice 3 weeks to 2 months after viral injection.

### 4.3 In vivo two-photon calcium imaging

For imaging experiments, the animal was fitted with a head bar, and head-fixed while resting on a rotating treadmill. The animal was awake and free to move on the treadmill, but not engaged in any conditioned behavior. Two-photon imaging was performed on a custom-built microscope with a 16×, 0.8 NA, 3 mm WD objective (Nikon) and controlled by custom software written in Labview (National Instruments). A Ti:Sapphire laser with mode-locking technique (Spectra-Physics Mai Tai HP DeepSee) was scanned by galvanometers (Cambridge). GCaMP6f was excited at 920 nm and laser power at the sample plane was typically 20 − 80 mW. A × 600 µm 600 µm field of view was scanned at 4.8 Hz as a series of × 250 pixel 250 pixel images and the imaging depth was up to 350 µm. Emitted light was collected with a T600/200dcrb dichroic (Chroma), passed through a HQ575/250m-2p bandpass filter (Chroma), and detected by a photomultiplier tube (R3896, Hamamatsu). Artifacts of the strobed stimulus (see below) were eliminated by discarding 8 pixels on either end of each line. The animal’s locomotion on the treadmill and its pupil positions were recorded and synchronized to the image acquisition. The head-fixed animal performs only rare eye movements and locomotion (Li et al., 2020).

### 4.4 Visual stimulation

An LCD screen with LED backlight was placed 18 cm away from the mouse’s right eye. The center of the monitor was at 95° azimuth and 25° elevation in the laboratory frame, and the monitor covered a visual field of 106° × 79°. The visual angle that covers the receptive fields of recorded neurons is 60°-140° azimuth and 0°-50° elevation (Figure 10). The monitor’s LED illuminator was strobed for 12 µs at the end of each laser scan line to minimize interference of the stimulus with fluorescence detection. The monitor was gamma-corrected. For measuring the functional properties, we presented six types of visual stimuli. (1) A full-field moving black bar (5° width at 50°/s) in 12 directions to measure the orientation selectivity and direction selectivity. The sequence of directions was pseudo-randomized. (2) An expanding black disc (diameter 2° to 60° at a speed of 60°/s, stationary at 60° for 0.25 s, followed by grey background for 2 s) and a receding white disc (60° to 2° at a speed of 60°/s, other parameters same as expanding disc) to measure looming-related responses. (3) Sparse (one at a time) 5° × 5° flashing squares (11 × 11 squares, 1 s black or white + 1 s grey) to map the receptive field (RF); (4) A 10° × 10° square modulated by a “chirp” in frequency or amplitude (3 s black + 3 s white + 3 s black + 3 s grey + 8 s frequency modulation (2^*−*1:3^ Hz) + 3 s grey + 8 s amplitude modulation (0 : 1) + 3 s grey + 3 s black) centered on the RF to measure temporal properties (Baden et al., 2016); (5) A 10° × 10° square flashing blue or green (1 s black + 3 s blue + 4s black + 3 s green + 3 s black) centered on the RF to measure the color preference; (6) A flashing disc (2 s black + 2 s grey) with different size (2°, 4°, 8°, 16°, 32°) centered on the RF to measure the size tuning. All stimuli were displayed for 10 repetitions. Stimuli of types 4, 5, and 6 were also repeated identically at locations in a 3 × 3 array shifted by 10°. This effectively covered the visual field recorded during a given imaging session. For each neuron, we based the analysis on the stimulus closest to its receptive field center.

### 4.5 Analysis of calcium responses

#### 4.5.1 Measurement of calcium responses

Brain motion during imaging was corrected using SIMA (Kaifosh et al., 2014) or NoRMCorre (Pnevmatikakis and Giovannucci, 2017). Regions of interest (ROIs) were drawn manually using Cell Magic Wand Tool (ImageJ) and fitted with an ellipse in MATLAB. Fluorescence traces of each ROI were extracted after estimating and removing contamination from surrounding neuropil signals as described previously (Feinberg and Meister, 2015; Li et al., 2020; Göbel and Helmchen, 2007; Kerlin et al., 2010). The true fluorescence signal of a neuron is *F*_true_ = *F*_raw_ − (*r F*_neuropil_), where *r* is the out-of-focus neuropil contamination factor and the estimated value for our setup is ∼ 0.7. Slow baseline fluctuations were removed by subtracting the eighth percentile value from a 15 s window centered on each frame (Dombeck et al., 2007).

For any given stimulus, the response of a neuron was defined by the fluorescence trace in its ROI during the stimulus period:

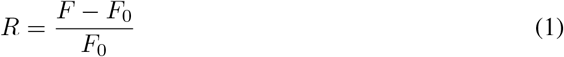

where *F* is the instantaneous fluorescence intensity and *F*_0_ is the mean fluorescence intensity without visual stimulation (grey screen).

Two criteria were applied to interpret ROIs as neurons: 1) The size of the ROI was limited to 10-20 µm to match the size of a neuron; 2) The response from the ROI had to pass a signal-to-noise ratio (SNR) of 0.35 (Baden et al., 2016).

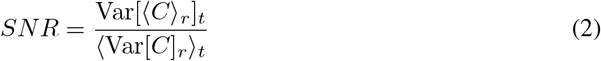

Where *C* is the *N*_*t*_ (time samples) × *N*_*r*_ (stimulus repetitions) response matrix, *t* = 1, …, *N*_*t*_ and *r* = 1, …, *N*_*r, r*_ and _*t*_ are the means over repetitions or time respectively, and Var[.]_*r*_ and Var[.]_*t*_ are the corresponding variances. All ROIs meeting these criteria were selected for further analysis, yielding a total of 3,414 neurons.

#### 4.5.2 Quantification of functional properties

The functional properties introduced in Figure 2 are defined as follows:

The response to motion (RtM) is the response value during moving-bar stimuli with the largest absolute value. For neurons suppressed by motion this will be negative.

To quantify the tuning of a neuron to motion directions, we calculated the direction selectivity index (DSI) as the normalized amplitude of the response-weighted vector sum of all directions:

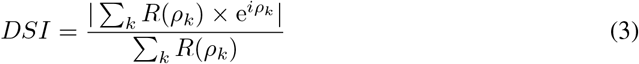

Where *ρ*_*k*_ is the *k*^th^ direction in radians and *R*(*ρ*_*k*_) is the peak response at that direction.

To quantify the orientation tuning, we calculated the orientation selectivity index (OSI) as the normalized amplitude of the response-weighted vector sum of all orientations:

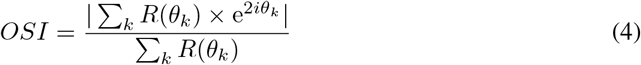

Where *θ*_*k*_ is the *k*^th^ orientation in radians and *R*(*θ*_*k*_) is the peak response at that orientation.

To quantify the habituation to the expanding dark disc, we calculated the habituation index (HI):

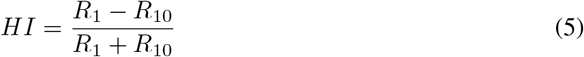

where *R*_1_ and *R*_10_ are the peak response to the first and the tenth looming stimulus respectively.

To quantify the selectivity to the expanding black disc over the receding white disc, we calculated the looming selectivity index (LSI):

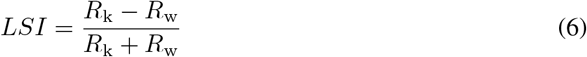

where *R*_k_ is the peak response to the black expanding disc and *R*_w_ is the peak response to the white receding disc.

To quantify the selectivity to moving stimuli over the flashing stimuli, we calculated the motion selectivity index (MSI):

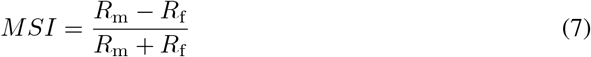

where *R*_m_ is the peak response to the moving bar at the preferred direction and *R*_f_ is the peak response to the flashing chirp stimulus.

To quantify the selectivity to On/Off contrast, we calculated the contrast selectivity index (CSI):

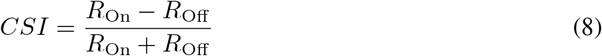

where *R*_On_ is the peak response to the flashing white square and *R*_Off_ is the peak response to the flashing black square.

To quantify whether neurons show transient or sustained responses to flash stimuli, we calculated the peak-final selectivity index (PFSI):

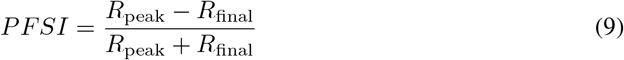

where *R*_peak_ is the peak response to the flashing white/black square that elicited larger responses, and *R*_final_ is the final response to that stimulus.

The quantify the selectivity to the flash frequency, we calculated the frequency selectivity index (FSI):

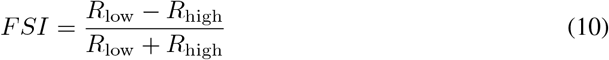

where *R*_low_ is the peak response in the first 3 seconds to the flashing frequency modulation, and *R*_high_ is the peak response in the last 2 seconds to the frequency modulation.

We measured the response after frequency modulation (RaFM) as the difference between the response amplitude at 1.6 s after the stop of the frequency modulation and the baseline. Similarly, the response after amplitude modulation (RaAM) was measured as the difference between the response amplitude at 1.6 s after the stop of the amplitude modulation and the baseline.

The best stimulation size (BSS) was defined the size of flashing black disc that elicited the largest responses.

To quantify the surrounding suppression, we calculated the surrounding suppression index (SSI):

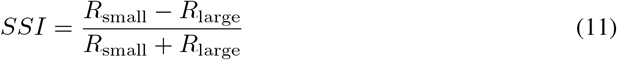

where *R*_small_ is the peak response to the flashing black disc with a diameter of 2 degrees, and *R*_large_ is the peak response to the flashing black disc with a diameter of 32 degrees.

To quantify the color preference, we calculated the blue-green index (BGI):

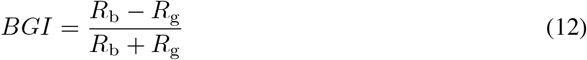

where *R*_b_ is the response to the flashing blue stimulus and *R*_g_ is the response to the flashing green stimulus.

To quantify the receptive field size (RFS), the calcium responses at 11 × 11 locations are fitted with a 2-D Gaussian function (Equation 13),

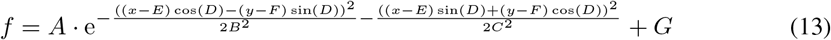

The RF size is defined as the area at the tenth of maximum, which equals *π* 2 ln 10 *BC*. We omitted analysis of the RF if the coefficient of determination for this fit was below 0.5 (Figure 6I). The RF size of neurons with the coefficient of determination larger than 0.5 is shown in Figures 2-4.

#### 4.5.3 Construction of the feature matrix

To construct the feature matrix, we first extracted features from neuronal responses to visual stimuli, including moving bars (MB), expanding black and receding white disc (EBD and RWD), chirp, color, flashed black discs with different sizes (FDDS). Neuronal responses to MB, EBD, and RWD were aligned to the peak or the trough to remove the effect of RF position. We focused on neurons which respond robustly (SNR>0.35) to at least one stimulus. For each neuron and each stimulus, the response was normalized to [0,1] and features were then extracted with sparse principal components analysis (spca) (Mairal et al., 2009), as implemented in the scikit-learn package (equation 14).

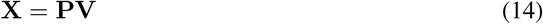

where **X** (*N*_neuron_ × *N*_time_) is the matrix of responses to a visual stimulus, **P**(*N*_neuron_ × *N*_feature_) is the matrix of principal components. **V**(*N*_feature_ × *N*_time_) is regularized so that only sparse values in each row are non-zero, and **VV**^*T*^ ≈ **I**.

We extracted 6 features from the responses to MB at the preferred direction, 6 features from the responses to EBD and RWD, 20 features from the chirp, 8 features from color stimuli, and 10 features from FDDS. These extracted features were combined with HI, DSI, OSI, and MSI to make the feature matrix. Each feature was normalized so that the mean is 0 and the standard deviation is 1.

#### 4.5.4 Clustering of the feature matrix

We used a Gaussian mixture model (GMM) to fit the distribution of neurons in the space of features.

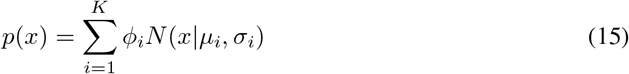

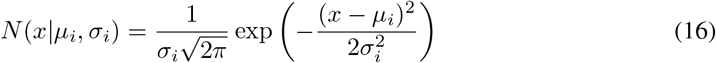

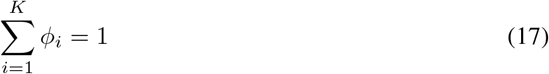

where *p*(*x*) is the probability density of the feature vector *x, K* is the number of component Gaussian functions, and *φ*_*i*_ is the weight for *i*^th^ Gaussian function *N* (*x* | *µ*_*i*_, *σ*_*i*_) in the feature space. We optimized the parameters using the EM algorithm (sklearn.mixture.GaussianMixture in the package scikit-learn). We varied the number of components from 2 to 50 and evaluated the quality with the Bayesian information criterion (BIC) (Kass and Raftery, 1995):

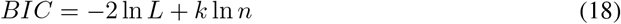

where *L* = *p*(*x* | *θ, M*), is the maximized likelihood of model *M, x* is the observed data, *θ* are the parameters that maximize the likelihood, *k* is the number of parameters in the model, *n* is the number of neurons. For each putative number of components (2 to 50), we performed the EM fit starting from 1000 random initial states, and chose the fit with the smallest BIC. This minimal BIC is plotted against the number of components in Figure 2B.

To assess the robustness of classification by the EM algorithm, we measured the probability that a pair of cells is classified into the same cluster in fits starting from different initial states, and calculated the co-association matrix (Fred and Jain, 2005):

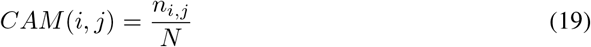

where *n*_*i,j*_ is the number of times that the pair (*i, j*) is assigned to the same cluster in *N* attempts. The between-cluster rate is defined as the cluster-wise average of the co-association matrix.

To evaluate the stability of clusters, we applied sub-sampling analysis (Hennig, 2007). We randomly sub-sampled 90% of the dataset 1000 times and fitted the subset with a GMM using the best cluster number determined from the full dataset. For each original cluster, we calculated its Jaccard similarity coefficient (JSC) with the subsets,

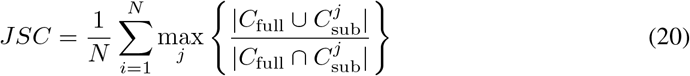

where *N* is the number of subsets, *C*_full_ is the cluster in the full dataset, and 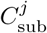 is the *j*th cluster in one subset. Clusters with JSC below 0.5 were considered unstable. The unstable cluster was merged with the cluster that has the highest between-cluster rate if it is *>* 35% (Gouwens et al., 2019); otherwise, it was marked in the figure.

We also explored clustering of cell types based on subsets of the visual stimuli, for example only the chirp sequence or the moving bars. The same BIC criterion identified more clusters with the restricted stimulus sets than with the full complement (Figure 6B).

#### 4.5.5 Relative selectivity index

Relative selectivity index (RSI) is defined as the difference of functional property between one type and a reference number.

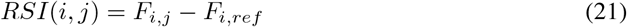

where *F*_*i,j*_ is functional property *i* of type *j* and *F*_*i,ref*_ is the reference of functional property *i*. The functional properties are response to motion (RtM), motion selectivity index (MSI), direction selectivity index (DSI), orientation selectivity index (OSI), habituation index (HI), looming selectivity index (LSI), contrast selectivity index (CSI), peak-final selectivity index (PFSI), response after frequency modulation (RaFM), response after amplitude modulation (RaAM), best stimulation size (BSS), surrounding suppression index (SSI), blue-green selectivity index (BGSI), receptive field size (RFS). The reference numbers are RtM: 0, DSI: 0.15, OSI: 0.15, HI: 0, LSI: 0, MSI: 0, CSI: 0, PFSI: 0.5, FSI: 0, RaFM: 0, RaAM: 0, BSS: 2^3^, SSI: 0, BGI: 0, RFS: 10^2.1^.

#### 4.5.6 Analysis of the anatomical arrangement of functional cell types

For the results on anatomical arrangement (Figure 3), only recording sessions with >5 neurons in a field of view were included. The density recovery profile (DRP) plots the probability per unit area of finding a cell as a function of distance from a cell of the same type (Rodieck, 1991). We first defined the region of interest (ROI) as the convex hull of all neurons in an image. Within this ROI, we measured the distances from each reference cell to all of the other cells and histogrammed those, which yields

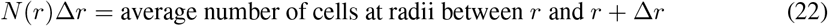

Then we measured the average area *A*(*r*)Δ*r* at distance between *r* and *r* + Δ*r* from any reference point in the window.

Finally the DRP was calculated as

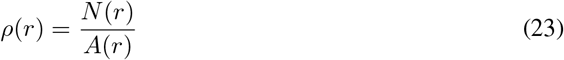

The density ratio (DR) for each type is calculated as

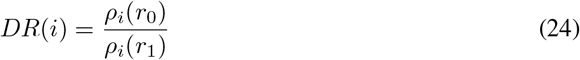

where *ρ*_*i*_(*r*_0_) is the mean density of functional type *i* within 0.5 average RF size of that type, and *ρ*_*i*_(*r*_1_) is the mean density in the annulus spanning 0.5-1 RF size. To connect anatomical distance in the SC with angular distance in the visual field we assumed that 1 mm corresponds to 88 degrees (Dräger and Hubel, 1976).

To quantify how the functional type of one neuron is related to the functional types of its neighbors, we calculated the density of different types

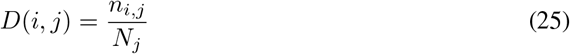

where *n*_*i,j*_ is number of neurons of functional type *j* within a certain distance to a neuron of type *i*, and *N*_*j*_ is the number of neurons of functional type *j* in the same area if neurons were uniformly distributed.

To quantify the relationship between two types of neurons, we calculated their normalized distance (ND) for each image

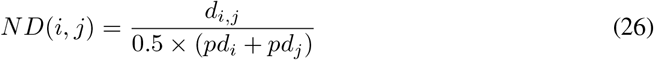

where *d*_*i,j*_ is the euclidean distance between the center of types *i* and *j*, and *pd*_*i*_ is the mean pairwise distance between neurons of type *i*.

To quantify the significance of the separation between two types, we shuffled labels for all neurons in these two clusters and calculated the p-value with bootstrap analysis to test whether the two types are significantly separated. If the maximum p-value of all images that have ≥ 10 neurons for both types is ≤ 0.01, these two types are significantly separated.

#### 4.5.7 Retina-SC transformation

For each type of SC neurons, we asked whether its responses can be explained by superposition of a small number of retinal ganglion cell types. Given the known responses of RGC types to these same stimuli (Baden et al., 2016) we approximated the response of each SC type as a weighted combination of RGC responses.

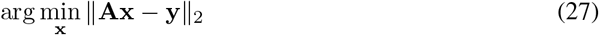

where **y** is the response vector of the SC type, **x** are the response vectors to the same stimuli for all types of RGCs, and **A** is the desired set of weights. The prediction error is quantified as

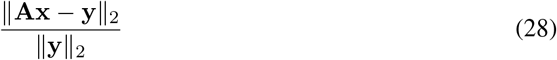

## 5 Supplement

Here we report details related to the Results and Methods sections.

**Figure 6:**
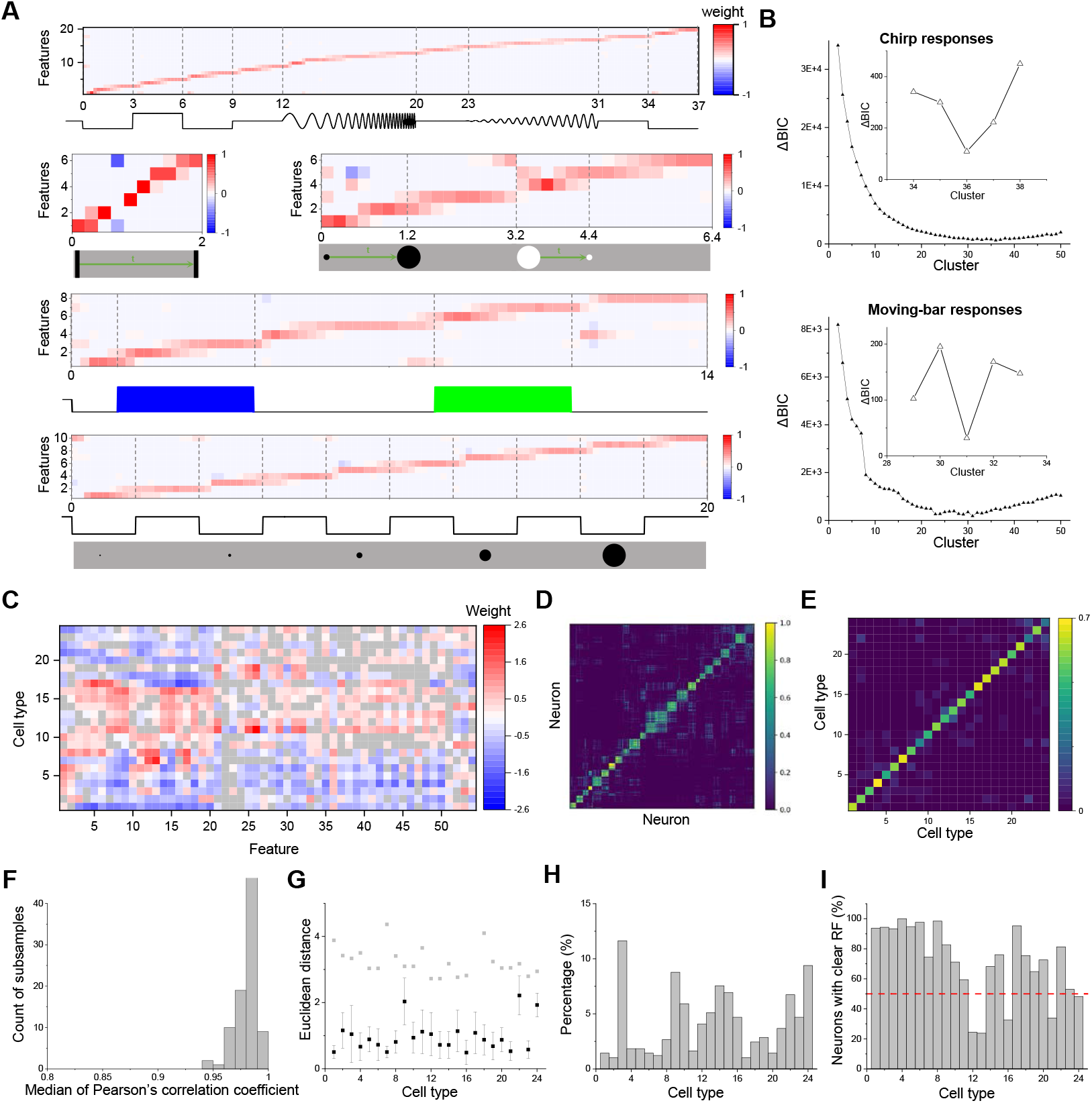
Clustering and validating (related to Figure 2). **A**. Temporal features were extracted from responses to five visual stimuli. Horizontal axis: time in s. **B**. Relative Bayesian information criterion (Δ BIC) for Gaussian mixture models with different numbers of clusters based on neuronal responses to chirp or moving-bar stimuli. **C**. Weight of 54 features in 24 cell types. Grey color indicates weights that are not significantly different from 0 (p>=0.05). **D**. Cell-wise co-association matrix (see Methods). Color bar indicates co-clustering fraction. **E**. Between-cluster rate, which is the cluster-wise average of the co-association matrix. **F**. Histogram of median correlation coefficients between the original clusters and clusters identified on 100 subsets. **G**. Black symbols indicate the Euclidean distance between original clusters and clusters identified on the subsets. Grey symbols indicate the maximal correlation coefficients between the original cluster and other clusters. **H**. Percentage of different functional types for neurons recorded in the SC of wild-type mice. **I**. Percentage of neurons that show clear RFs to flash stimuli for each cell type.

**Figure 7:**
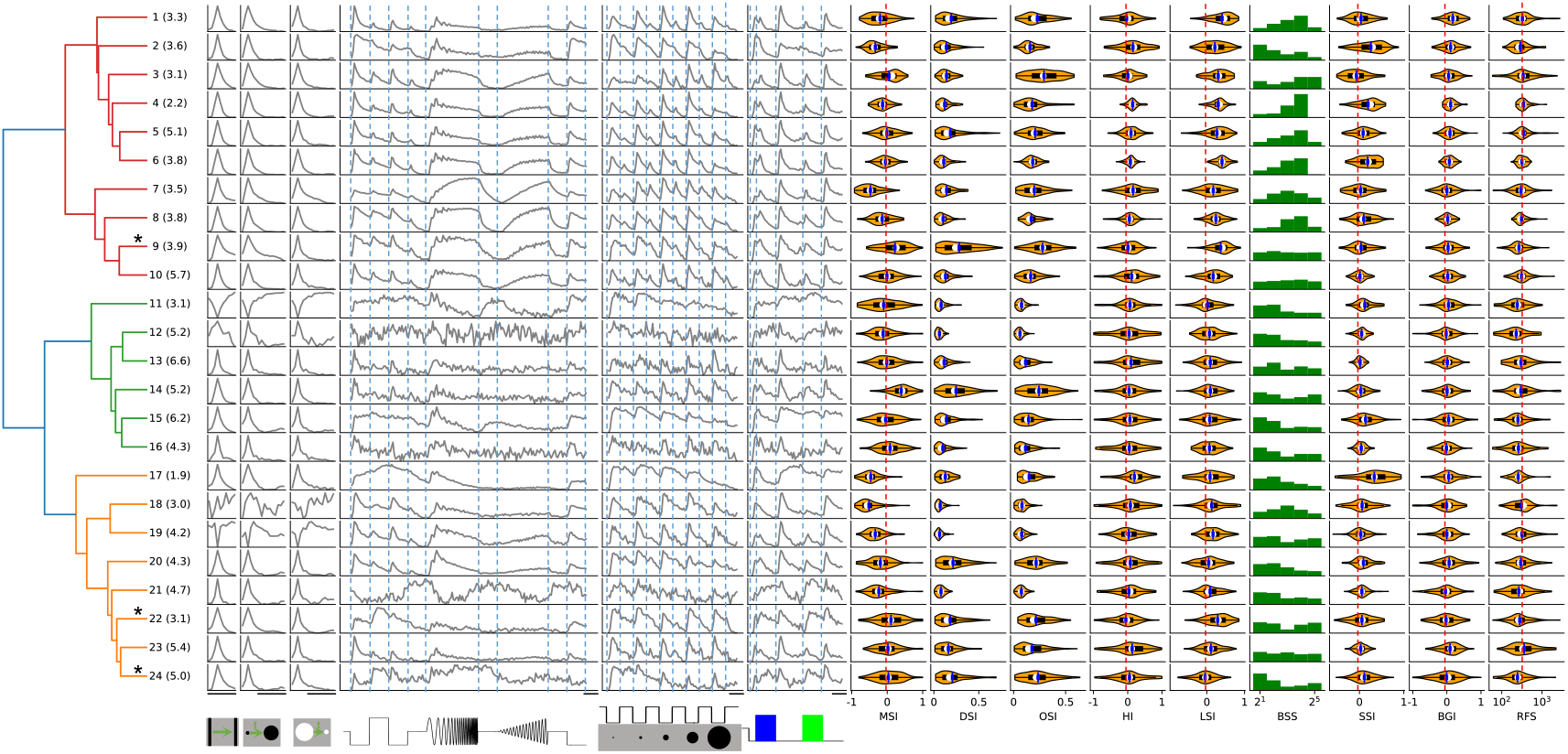
Dendrogram of 24 clusters showing normalized temporal profiles and functional properties (related to Figure 2). Numbers in the parentheses indicate percentage of each type. Display as in Figure 2A; scale bars: 2 s. Star marks the unstable cluster with JSC*<* 0.5.

**Figure 8:**
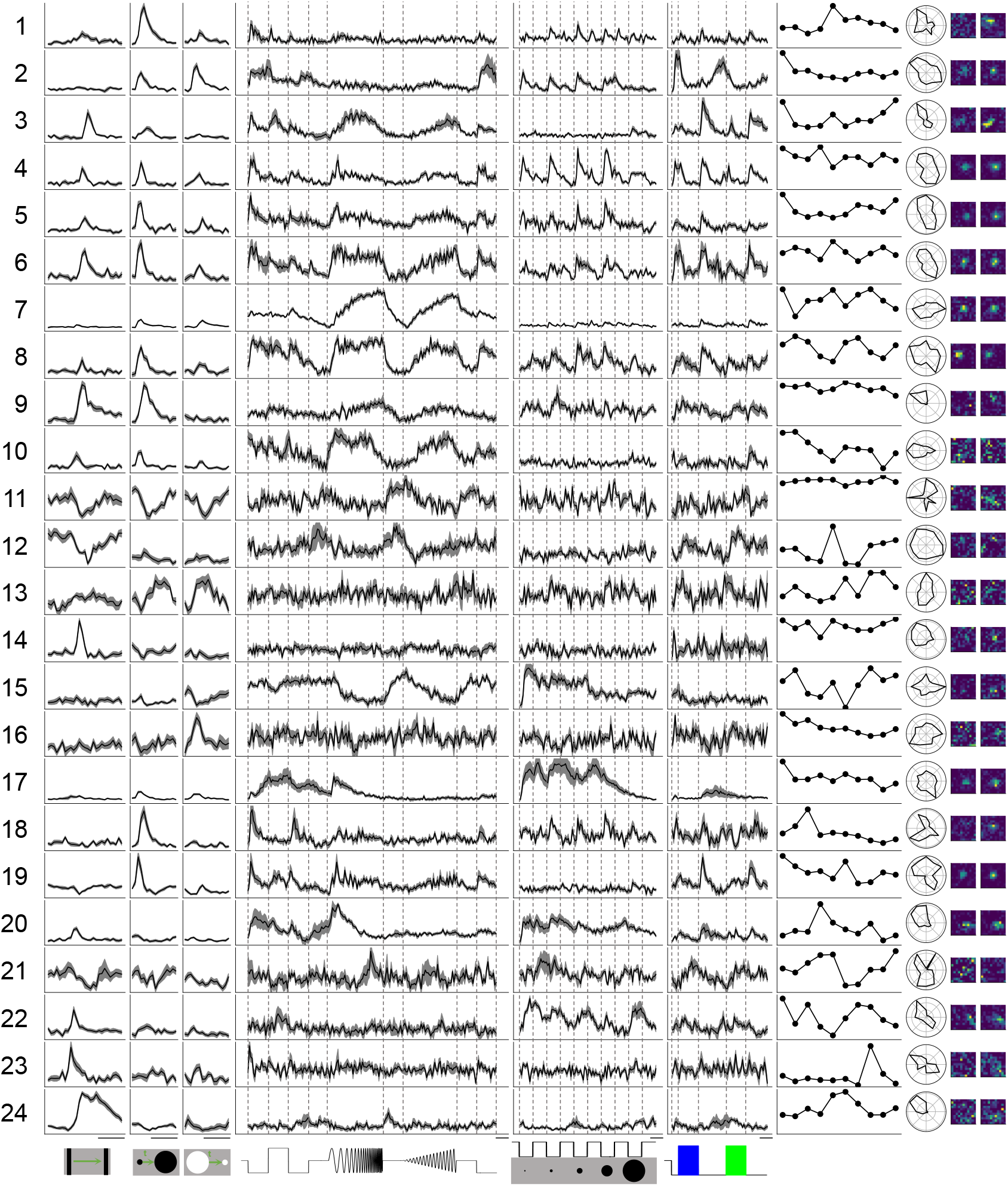
Example responses of each type (related to Figure 2). Display as in Figure 1D; scale bars: 2 s.

**Figure 9:**
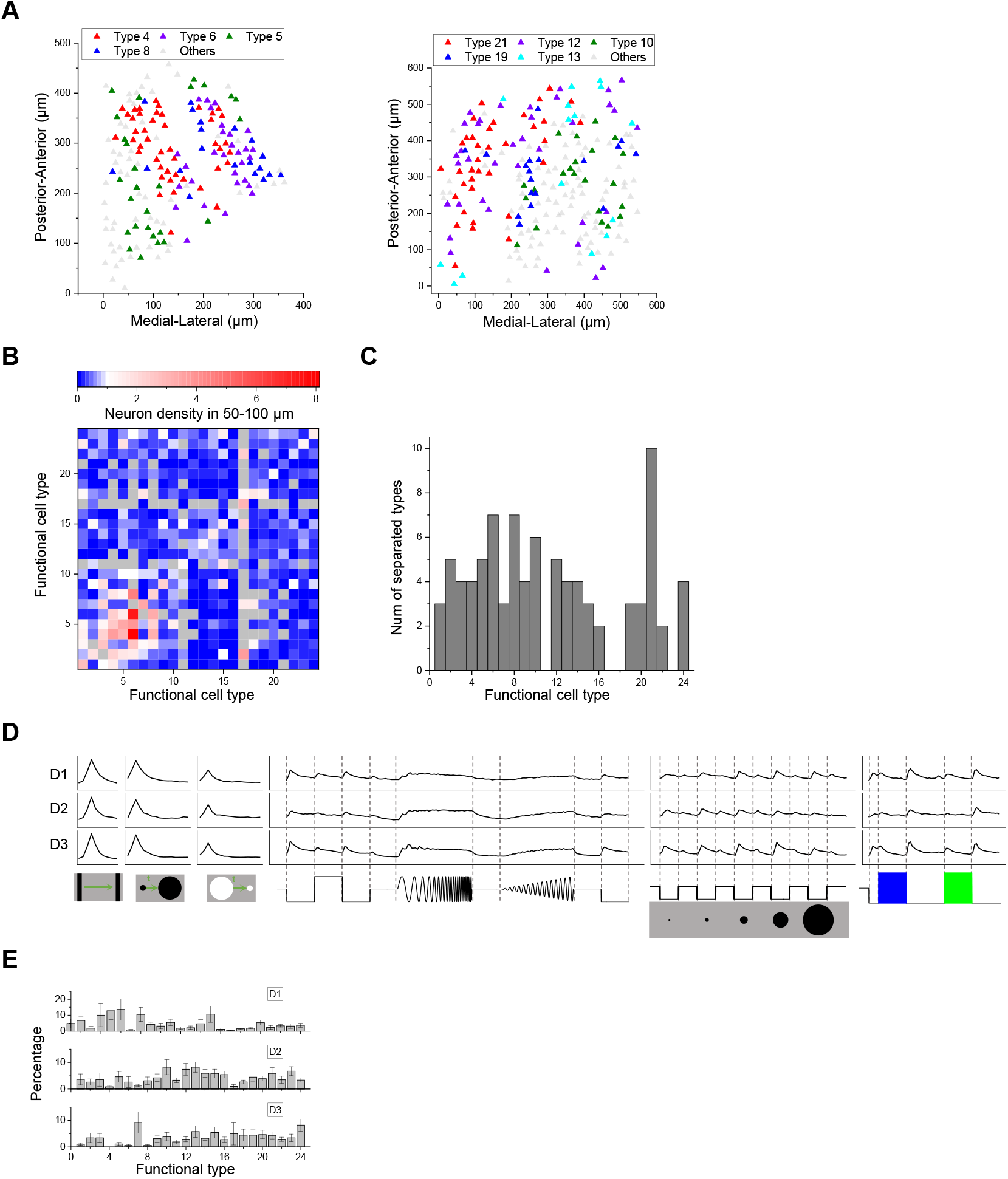
Anatomical organization of functional types (related to Figure 3). **A**. Anatomical locations of different types of neurons in two examples. **B**. Density of neurons in different functional types (columns) in 50-100 *µ*m of a given neuron whose type is indicated by the row. Grey color indicates insufficient data for those pairs. **C**. Number of types that are significantly separated from each type. **D**. Temporal response across depth. Display as in Figure 2A. **E**. The percentage of functional cell types across depth. Error bars denote SEM.

**Figure 10:**
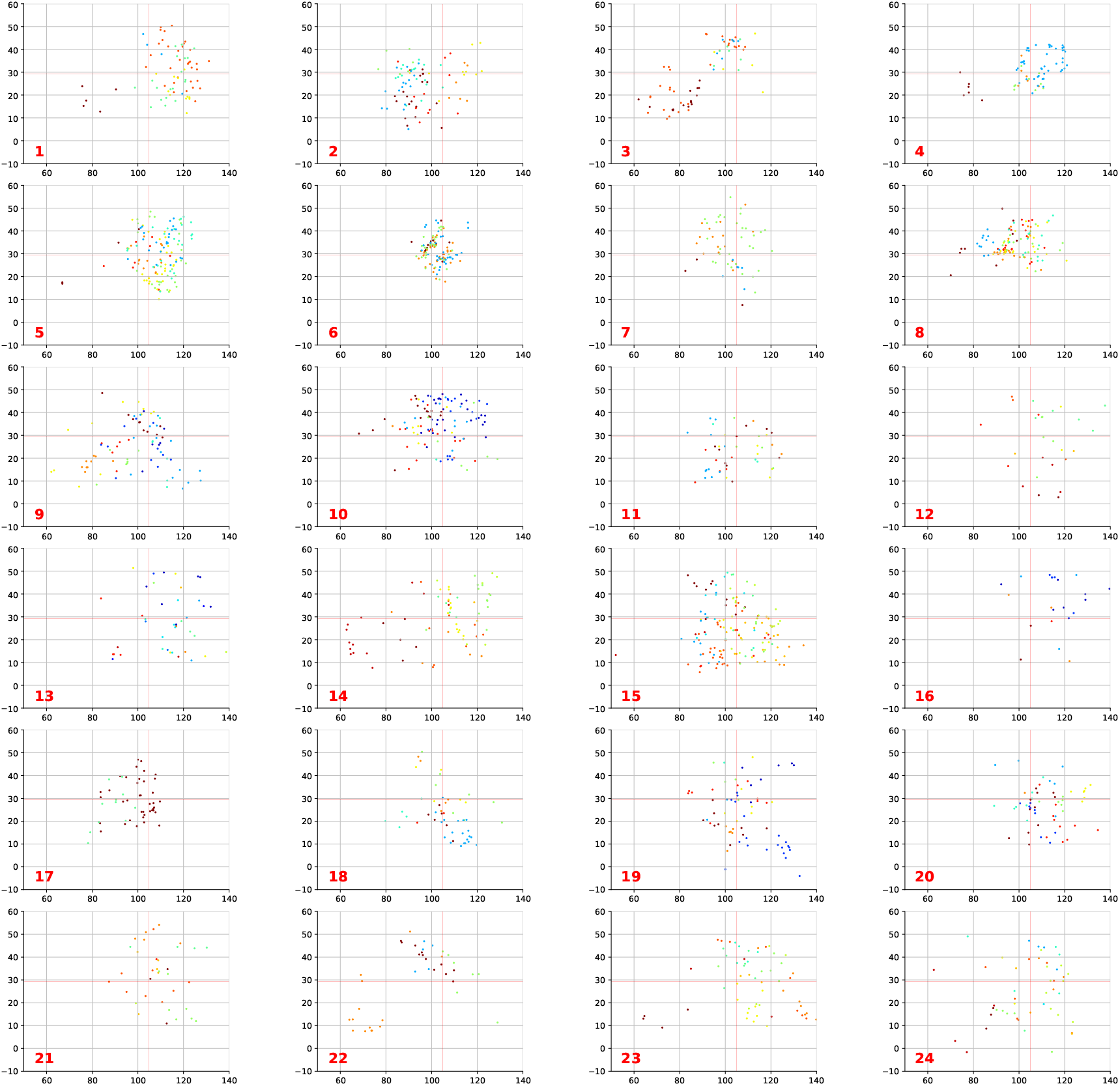
Receptive field center positions for each functional type (related to Figure 3). Cell types are indicated by the red number at the corner. Color indicates different experiments. Axes represent azimuth (horizontal) and elevation (vertical) in degrees.

**Figure 11:**
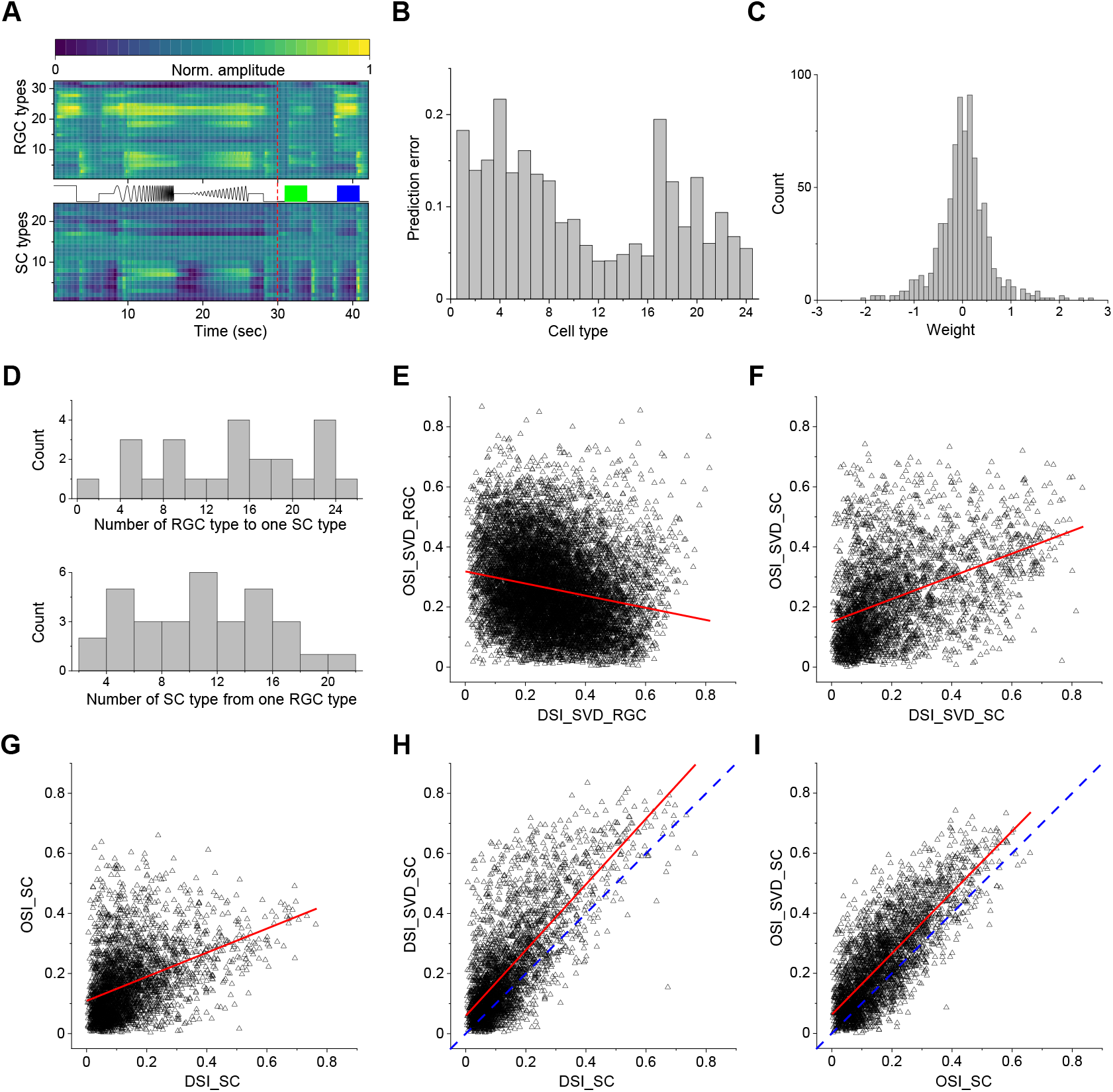
Comparisons between SC neurons and RGCs (related to Figure 5). **A**. Visual responses of RGC types (Baden et al., 2016) and SC types. **B**. Prediction error from RGC types to SC types (Eqn 28). **C**. Histogram of the weights of RGC types to SC types in Figure 5A. **D**. If one only considers weights | *w* | *>* 0.3, this histograms the number of RGC types contributing to one SC type, and vice versa the number of SC types with contributions from one RGC type. **E**. Plot of OSI vs DSI among RGCs, as reported in (Baden et al., 2016). **F**. Plot of OSI vs DSI for SC neurons, calculated from the present study by the same SVD algorithm used in (Baden et al., 2016). **G**. Plot of OSI vs DSI for SC neurons, using the simpler definition from the present study. **H**. Plot of DSI for SC neurons, computed by the SVD algorithm (Baden et al., 2016) versus the present definition. Note the close correspondence. **I**. As panel (G) for OSI. This analysis shows that the comparison between the retina results in (Baden et al., 2016) and SC results in the present study does not suffer from different analysis methods.

## Funding

M.M. was supported by grants from NIH (R01 NS111477) and from the Simons Foundation (543015SPI). Y.-t.L was supported by a grant from the NEI (K99EY028640) and a Helen Hay Whitney Postdoctoral Fellowship.

## Author contributions

Y.-t.L designed the study, performed all experiments, interpreted results, and wrote the manuscript. M.M. helped design the study, interpret results, and write the manuscript.

## Competing interests

The authors declare no competing interests.

## Data and code availability

Data and code will be available in a public repository following acceptance of the manuscript.

## Notes

### Competing Interest Statement

The authors have declared no competing interest.

### Summary of Updates

Both manuscript and figures are updated.

## References

Ahmadlou, M. and Heimel, J. A. (2015). Preference for concentric orientations in the mouse superior colliculus. Nature Communications, 6:6773.

Baden, T., Berens, P., Franke, K., Román Rosón, M., Bethge, M., and Euler, T. (2016). The functional diversity of retinal ganglion cells in the mouse. Nature, 529(7586):345–350.

Basso, M. A. and May, P. J. (2017). Circuits for Action and Cognition: A View from the Superior Colliculus. Annual Review of Vision Science, 3(1):197–226.

Blasdel, G. and Campbell, D. (2001). Functional Retinotopy of Monkey Visual Cortex. Journal of Neuroscience, 21(20):8286–8301.

Bleckert, A., Schwartz, G. W., Turner, M. H., Rieke, F., and Wong, R. O. L. (2014). Visual Space Is Represented by Nonmatching Topographies of Distinct Mouse Retinal Ganglion Cell Types. Current Biology, 24(3):310–315.

Byun, H., Kwon, S., Ahn, H.-J., Liu, H., Forrest, D., Demb, J. B., and Kim, I.-J. (2016). Molecular features distinguish ten neuronal types in the mouse superficial superior colliculus. Journal of Comparative Neurology, 524(11):2300–2321. _eprint: https://onlinelibrary.wiley.com/doi/pdf/10.1002/cne.23952.

Chandrasekaran, A. R., Shah, R. D., and Crair, M. C. (2007). Developmental Homeostasis of Mouse Retinocollicular Synapses. Journal of Neuroscience, 27(7):1746–1755.

Chen, H., Savier, E. L., DePiero, V. J., and Cang, J. (2021). Lack of Evidence for Stereotypical Direction Columns in the Mouse Superior Colliculus. Journal of Neuroscience, 41(3):461–473. Publisher: Society for Neuroscience Section: Research Articles.

De Franceschi, G. and Solomon, S. G. (2018). Visual response properties of neurons in the superficial layers of the superior colliculus of awake mouse. The Journal of Physiology, 596(24):6307–6332.

de Malmazet, D., Kühn, N. K., and Farrow, K. (2018). Retinotopic Separation of Nasal and Temporal Motion Selectivity in the Mouse Superior Colliculus. Current Biology, 28(18):2961–2969.e4.

Dombeck, D. A., Khabbaz, A. N., Collman, F., Adelman, T. L., and Tank, D. W. (2007). Imaging Large-Scale Neural Activity with Cellular Resolution in Awake, Mobile Mice. Neuron, 56(1):43–57.

Dräger, U. C. and Hubel, D. H. (1975). Responses to visual stimulation and relationship between visual, auditory, and somatosensory inputs in mouse superior colliculus. Journal of Neurophysiology, 38(3):690–713.

Dräger, U. C. and Hubel, D. H. (1976). Topography of visual and somatosensory projections to mouse superior colliculus. Journal of Neurophysiology, 39(1):91–101.

Ellis, E. M., Gauvain, G., Sivyer, B., and Murphy, G. J. (2016). Shared and Distinct Retinal Input to the Mouse Superior Colliculus and Dorsal Lateral Geniculate Nucleus. Journal of Neurophysiology, page jn.00227.2016.

Feinberg, E. H. and Meister, M. (2015). Orientation columns in the mouse superior colliculus. Nature, 519(7542):229–232.

Fred, A. L. N. and Jain, A. K. (2005). Combining multiple clusterings using evidence accumulation. IEEE Transactions on Pattern Analysis and Machine Intelligence, 27(6):835–850. Conference Name: IEEE Transactions on Pattern Analysis and Machine Intelligence.

Gale, S. D. and Murphy, G. J. (2014). Distinct Representation and Distribution of Visual Information by Specific Cell Types in Mouse Superficial Superior Colliculus. The Journal of Neuroscience, 34(40):13458–13471.

Gerfen, C. R., Paletzki, R., and Heintz, N. (2013). GENSAT BAC Cre-Recombinase Driver Lines to Study the Functional Organization of Cerebral Cortical and Basal Ganglia Circuits. Neuron, 80(6):1368–1383. Publisher: Elsevier.

Gouwens, N. W., Sorensen, S. A., Berg, J., Lee, C., Jarsky, T., Ting, J., Sunkin, S. M., Feng, D., Anastassiou, C. A., Barkan, E., Bickley, K., Blesie, N., Braun, T., Brouner, K., Budzillo, A., Caldejon, S., Casper, T., Castelli, D., Chong, P., Crichton, K., Cuhaciyan, C., Daigle, T. L., Dalley, R., Dee, N., Desta, T., Ding, S.-L., Dingman, S., Doperalski, A., Dotson, N., Egdorf, T., Fisher, M., Frates, R. A. d., Garren, E., Garwood, M., Gary, A., Gaudreault, N., Godfrey, K., Gorham, M., Gu, H., Habel, C., Hadley, K., Harrington, J., Harris, J. A., Henry, A., Hill, D., Josephsen, S., Kebede, S., Kim, L., Kroll, M., Lee, B., Lemon, T., Link, K. E., Liu, X., Long, B., Mann, R., McGraw, M., Mihalas, S., Mukora, A., Murphy, G. J., Ng, L., Ngo, K., Nguyen, T. N., Nicovich, P. R., Oldre, A., Park, D., Parry, S., Perkins, J., Potekhina, L., Reid, D., Robertson, M., Sandman, D., Schroedter, M., Slaughterbeck, C., Soler-Llavina, G., Sulc, J., Szafer, A., Tasic, B., Taskin, N., Teeter, C., Thatra, N., Tung, H., Wakeman, W., Williams, G., Young, R., Zhou, Z., Farrell, C., Peng, H., Hawrylycz, M. J., Lein, E., Ng, L., Arkhipov, A., Bernard, A., Phillips, J. W., Zeng, H., and Koch, C. (2019). Classification of electrophysiological and morphological neuron types in the mouse visual cortex. Nature Neuroscience, 22(7):1182–1195.

Göbel, W. and Helmchen, F. (2007). In Vivo Calcium Imaging of Neural Network Function. Physiology, 22(6):358–365.

Harris, J. A., Hirokawa, K. E., Sorensen, S. A., Gu, H., Mills, M., Ng, L. L., Bohn, P., Mortrud, M., Ouellette, B., Kidney, J., Smith, K. A., Dang, C., Sunkin, S., Bernard, A., Oh, S. W., Madisen, L., and Zeng, H. (2014). Anatomical characterization of Cre driver mice for neural circuit mapping and manipulation. Frontiers in Neural Circuits, 8. Publisher: Frontiers.

Hennig, C. (2007). Cluster-wise assessment of cluster stability. Computational Statistics & Data Analysis, 52(1):258–271.

Horn, G. and Hill, R. M. (1966). Responsiveness to sensory stimulation of units in the superior colliculus and subjacent tectotegmental regions of the rabbit. Experimental Neurology, 14(2):199–223.

Hoy, J. L., Bishop, H. I., and Niell, C. M. (2019). Defined Cell Types in Superior Colliculus Make Distinct Contributions to Prey Capture Behavior in the Mouse. Current Biology, 29(23):4130– 4138.e5.

Inayat, S., Barchini, J., Chen, H., Feng, L., Liu, X., and Cang, J. (2015). Neurons in the Most Superficial Lamina of the Mouse Superior Colliculus Are Highly Selective for Stimulus Direction. The Journal of Neuroscience, 35(20):7992–8003.

Isa, T., Marquez-Legorreta, E., Grillner, S., and Scott, E. K. (2021). The tectum/superior colliculus as the vertebrate solution for spatial sensory integration and action. Current Biology, 31(11):R741–R762. Publisher: Elsevier.

Ito, S., Feldheim, D. A., and Litke, A. M. (2017). Segregation of Visual Response Properties in the Mouse Superior Colliculus and Their Modulation during Locomotion. Journal of Neuroscience, 37(35):8428–8443.

Kaifosh, P., Zaremba, J. D., Danielson, N. B., and Losonczy, A. (2014). SIMA: Python software for analysis of dynamic fluorescence imaging data. Frontiers in Neuroinformatics, 8(80):1–10.

Kass, R. E. and Raftery, A. E. (1995). Bayes Factors. Journal of the American Statistical Association, 90(430):773–795.

Kerlin, A. M., Andermann, M. L., Berezovskii, V. K., and Reid, R. C. (2010). Broadly Tuned Response Properties of Diverse Inhibitory Neuron Subtypes in Mouse Visual Cortex. Neuron, 67(5):858–871.

Langer, T. P. and Lund, R. D. (1974). The upper layers of the superior colliculus of the rat: A Golgi study. Journal of Comparative Neurology, 158(4):405–435.

Lee, K. H., Tran, A., Turan, Z., and Meister, M. (2020). The sifting of visual information in the superior colliculus. eLife, 9:e50678. Publisher: eLife Sciences Publications, Ltd.

Li, Y.-t., Turan, Z., and Meister, M. (2020). Functional Architecture of Motion Direction in the Mouse Superior Colliculus. Current Biology, 30(17):3304–3315.e4.

Mairal, J., Bach, F., Ponce, J., and Sapiro, G. (2009). Online dictionary learning for sparse coding. In Proceedings of the 26th Annual International Conference on Machine Learning, ICML’09, pages 689–696, Montreal, Quebec, Canada. Association for Computing Machinery.

Major, D. E., Luksch, H., and Karten, H. J. (2000). Bottlebrush dendritic endings and large dendritic fields: Motion-detecting neurons in the mammalian tectum. The Journal of Comparative Neurology, 423(2):243–260.

May, P. J. (2006). The mammalian superior colliculus: laminar structure and connections. In Büttner-Ennever, J. A., editor, Progress in Brain Research, volume 151 of Neuroanatomy of the Oculomotor System, pages 321–378. Elsevier.

Pnevmatikakis, E. A. and Giovannucci, A. (2017). NoRMCorre: An online algorithm for piecewise rigid motion correction of calcium imaging data. Journal of Neuroscience Methods, 291:83–94.

Reinhard, K., Li, C., Do, Q., Burke, E. G., Heynderickx, S., and Farrow, K. (2019). A projection specific logic to sampling visual inputs in mouse superior colliculus. eLife, 8:e50697.

Rodieck, R. W. (1991). The density recovery profile: A method for the analysis of points in the plane applicable to retinal studies. Visual Neuroscience, 6(2):95–111. Publisher: Cambridge University Press.

Roska, B. and Meister, M. (2014). The retina dissects the visual scene into distinct features. In Werner, J. S. and Chalupa, L. M., editors, The new visual neurosciences, pages 163–82. MIT Press, Cambridge, MA.

Roy, S., Jun, N. Y., Davis, E. L., Pearson, J., and Field, G. D. (2021). Inter-mosaic coordination of retinal receptive fields. Nature, pages 1–5. Publisher: Nature Publishing Group.

Sanes, J. R. and Masland, R. H. (2015). The Types of Retinal Ganglion Cells: Current Status and Implications for Neuronal Classification. Annual Review of Neuroscience, 38(1):221–246.

Savier, E. L., Chen, H., and Cang, J. (2019). Effects of Locomotion on Visual Responses in the Mouse Superior Colliculus. Journal of Neuroscience, 39(47):9360–9368. Publisher: Society for Neuroscience Section: Research Articles.

Shekhar, K., Lapan, S. W., Whitney, I. E., Tran, N. M., Macosko, E. Z., Kowalczyk, M., Adiconis, X., Levin, J. Z., Nemesh, J., Goldman, M., McCarroll, S. A., Cepko, C. L., Regev, A., and Sanes, J. R. (2016). Comprehensive Classification of Retinal Bipolar Neurons by Single-Cell Transcriptomics. Cell, 166(5):1308–1323.e30. Publisher: Elsevier.

Wang, L., Sarnaik, R., Rangarajan, K., Liu, X., and Cang, J. (2010). Visual Receptive Field Properties of Neurons in the Superficial Superior Colliculus of the Mouse. Journal of Neuroscience, 30(49):16573–16584.

Warwick, R. A., Kaushansky, N., Sarid, N., Golan, A., and Rivlin-Etzion, M. (2018). Inhomogeneous Encoding of the Visual Field in the Mouse Retina. Current Biology, in press(0).

Wassle, H., Peichl, L., and Boycott, B. B. (1981). Morphology and Topography of on- and off-Alpha Cells in the Cat Retina. Proceedings of the Royal Society of London. Series B, Biological Sciences, 212(1187):157–175.

Yan, W., Laboulaye, M. A., Tran, N. M., Whitney, I. E., Benhar, I., and Sanes, J. R. (2020). Mouse Retinal Cell Atlas: Molecular Identification of over Sixty Amacrine Cell Types. Journal of Neuroscience, 40(27):5177–5195. Publisher: Society for Neuroscience Section: Research Articles.

Zeng, H. and Sanes, J. R. (2017). Neuronal cell-type classification: challenges, opportunities and the path forward. Nature Reviews Neuroscience, 18(9):530–546. Number: 9 Publisher: Nature Publishing Group.

Zhang, Y., Kim, I.-J., Sanes, J. R., and Meister, M. (2012). The most numerous ganglion cell type of the mouse retina is a selective feature detector. Proceedings of the National Academy of Sciences, 109(36):E2391–E2398.

